# Mycoplasma immunoglobulin binding protein universally binds human antibodies, thereby reducing Fab arm flexibility and displacing antigens in immune complexes

**DOI:** 10.64898/2026.06.12.731821

**Authors:** Tereza Kadavá, Awital Bar Barroeta, Carla de Haas, Suzan H.M. Rooijakkers, Jürgen Strasser, Johannes Preiner, Maria Nordgren, Rolf Lood, Albert J.R. Heck

## Abstract

Mycoplasma immunoglobulin binding (MIB) protein plays a key role in immune evasion across several Mycoplasma species. MIB tightly interacts with the Fab domain of immunoglobulin G, inducing conformational changes that disrupt the antigen-binding site. Here, we reveal that MIB binds strongly and universally to all major isotypes and subclasses of human antibodies and can thus serve as a bait protein to efficiently deplete antibodies from serum. Our results demonstrate strong interactions between MIB and a diverse set of both recombinant monoclonal and endogenous polyclonal antibodies, the latter purified from serum or colostrum. All antibodies, including IgG and IgA monomers, J-chain coupled IgA dimers, and IgM pentamers, consistently bind two copies of MIB per antibody protomer. Using cross-linking mass spectrometry, we pinpoint that antibody–MIB interactions involve conserved regions of the variable and constant antibody domains, independent of isotype. Furthermore, we demonstrate that MIB not only disrupts the Fab–antigen interaction but also restricts Fab flexibility, as visualized by atomic force microscopy. Conversely, MIB does not affect Fc-mediated protein interactions, as exemplified by the successful reconstitution of a 36-subunit immune complex, (IgG)_6_-(MIB)_12_-C1q.

## Introduction

Antibodies are key players in humoral immunity, mediating the detection, neutralization, and clearance of bacterial and viral infections.^1^ Antibody binding to bacterial surfaces via the fragment antigen-binding (Fab) domain (Fig. 1) can readily neutralize the target. Additionally, the antibody fragment crystallizable (Fc) region (Fig. 1) mediates effector functions.^2^ The Fc interacts with receptors on immune cells or complement proteins to initiate phagocytosis, intracellular killing, and the complement cascade.^3,4^ Complement activation can lead to the formation of the membrane attack complex (MAC), which, in more susceptible Gram-negative bacteria, results in direct lysis.^5^ Human circulation contains three major antibody isotypes: IgG, IgA, and IgM (Fig. 1), and low-abundant IgD and IgE. Furthermore, subclasses are distinguished within IgG and IgA isotypes: IgG1, IgG2, IgG3, IgG4, IgA1, and IgA2.^2^ IgA and IgM isotypes can form oligomeric complexes.^6^ While circulatory IgA is mostly monomeric, secretory IgA is predominantly a dimeric complex consisting of two IgA monomers joined by a J-chain and bound to the secretory component pIgR.^7^ IgM predominantly forms J-chain coupled pentamers, associated with CD5L or secretory component when circulating in blood or secretory biofluids, respectively.^8–10^ The antibody isotype that drives the immune response determines the resulting immune response. While all main antibody isotypes aid pathogen clearance by inducing phagocytosis, the complement pathway is primarily activated either by hexamers of IgG1–IgG3 or by IgM.^1,11–15^

**Figure 1.**
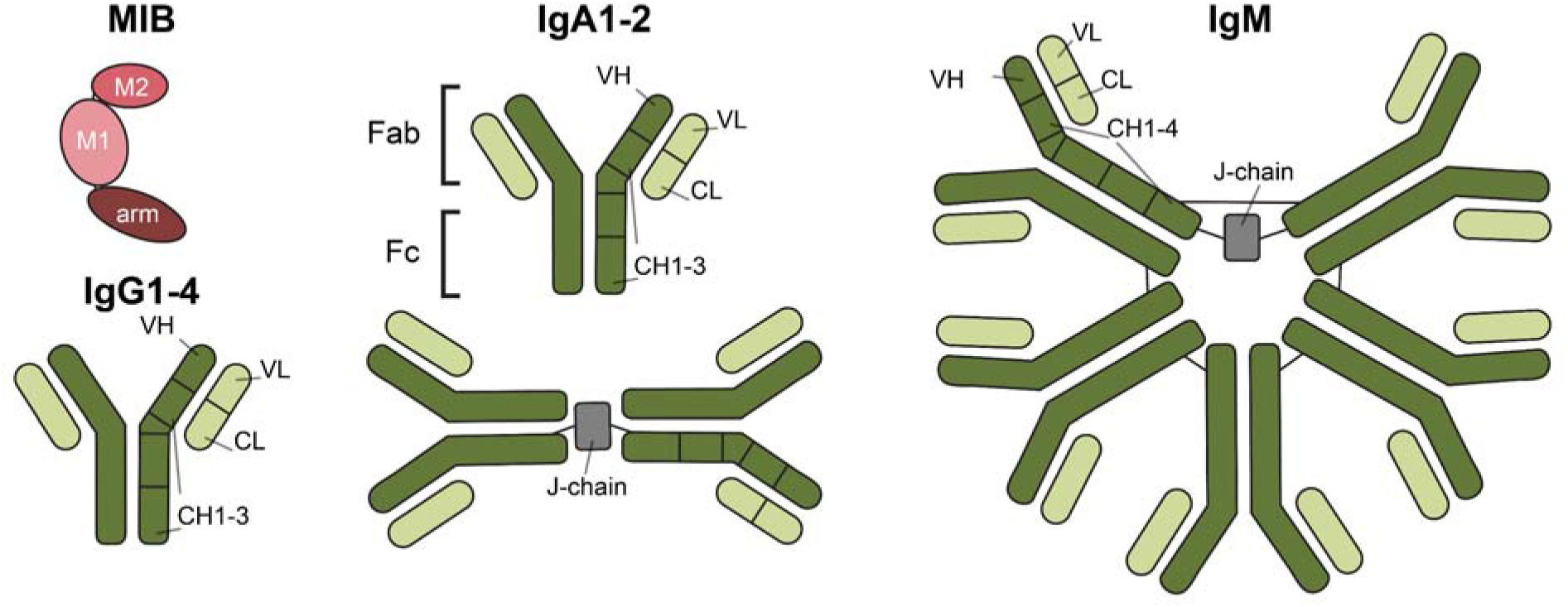
Schematic domain structures of MIB and human antibody (sub)classes. The structure of soluble MIB (red), which mediates antibody binding, consists of three domains: an N-terminal arm and the M1 and M2 domains.^28^ The antibody protomer structure resembles a Y-shape and is composed of two heavy chains (dark green) and two light chains (light green). Each antibody protomer contains two Fab domains that recognize and bind antigens. Antigen specificity is determined by the variable domains of the heavy (VH) and light (VL) chains. The constant (Fc) domain mediates antibody effector functions. Human circulation predominantly contains three isotypes: IgG, IgA, and IgM. IgG and IgA contain three constant heavy chain domains (CH), whereas IgM has four CH. Furthermore, IgA and IgM can exist as oligomeric structures bound to the J-chain. IgA mostly forms J-chain coupled dimers, while IgM J-chain coupled pentamers. These J-chain coupled oligomers can bind the secretory component of pIgR or CD5L in milk or serum, respectively.^1,6–8^

In response to the evolutionary pressure of the humoral immune system, bacteria have evolved various strategies to evade immune surveillance.^16,17^ These include the recruitment of complement regulators, the secretion of proteases that target components of the innate immune response, and the generation of antibody-binding proteins. Most bacterial antibody-binding proteins, such as staphylococcal protein A and streptococcal protein G, bind Fc and inhibit its effector functions. Conversely, mycoplasmas express the mycoplasma immunoglobulin binding (MIB) protein, which prevents initial pathogen recognition by targeting the Fab.^16,18–20^

Mycoplasmas are cell-wall-less bacteria that rely on their host for nutrient supply due to a drastic genome reduction that limits their metabolic capabilities.^21–24^ Despite their rudimentary nature, Mycoplasmas can infect a wide range of hosts. They primarily reside on mucosal surfaces lining respiratory, urogenital, and gastrointestinal tracts, mammary glands, eyes, and joints.^24^ Mycoplasma infections are persistent and generally cause chronic inflammatory diseases with low mortality. The chronicity of these infections can be attributed to the ability of mycoplasmas to effectively undermine the host’s immune system.^25,23,26^ MIB is part of a two-protein antibody–cleaving system known as the MIB–mycoplasma immunoglobulin protease (MIB–MIP) system, which forms a prevalent and elegant immune evasion strategy in animal-infecting Mycoplasma species.^20,26–28^

The MIB–MIP system interferes with the humoral immune system in two steps. First, MIB captures an antibody that targets bacterial surface proteins and then recruits the MIP protease. Next, MIP cleaves off a variable region on the heavy chain from the Fab domain, producing an antibody fragment that can no longer bind its cognate antigen.^28^ Notably, MIB alone can displace antibody–antigen interactions, in clear contrast to other antibody-binding proteins such as protein A and protein G. Upon binding to an antibody, MIB encircles the antibody light chain, inducing conformational changes.^28^ These conformational changes not only accommodate MIP binding but also disrupt the complementarity-determining regions, thereby preventing antigen binding and displacing antigen from existing antibody–antigen complexes.^28^ All this knowledge stems from research that was predominantly performed with polyclonal IgG-derived Fabs, and MIB interactions with other isotypes and polymeric assemblies have not been characterized extensively.

Here, we systematically characterized antibody–MIB binding using a diverse selection of recombinant monoclonal antibodies and polyclonal endogenous antibodies. Our data suggest that MIB binds antibodies strongly and universally, regardless of isotype or oligomeric composition. In all cases, each studied antibody bound to two MIB molecules per protomer. Furthermore, our results demonstrate that MIB interaction with an antibody not only affects the Fab structure but also substantially reduces its flexibility. While this affects the antigen-recognizing function of the Fab domain, the effector functions of the Fc domain remain largely intact. This is highlighted by the incorporation of MIB into the IgG1-C1q immune complex, as evidenced by the successful reconstitution of an (IgG1)_6_-(MIB)_12_-C1q immune complex. In summary, our data provide a comprehensive overview of MIB binding and underscore its potential as a bait protein for affinity purification and antibody depletion from serum.

## Results

The major aim of this work was to characterize MIB and its interactions across human antibody isotypes. Thus far, MIB interactions have been characterized with polyclonal goat IgG^20,28^. These results have positioned MIB as a promising biotechnological tool for targeting all human antibodies, which we aimed to further exploit.

### MIB is a specific pan-antibody binder

First, to elucidate MIB’s binding characteristics, affinities, and potential off-target interactors, we performed an affinity pull-down from human serum using MIB as a bait. We used recombinant MIB immobilized on NHS-activated agarose and collected flow-through (FT) and MIB-captured eluate. The protein abundances in each fraction, as well as in a serum control, were then analyzed by DIA LC-MS/MS based quantitative proteomics (Fig. 2A–B, File S1). The pooled serum control contained an expected ∼17% of antibodies.^29^ Quantitative proteomics revealed that all antibodies were effectively depleted in the FT, accounting for only 1.6% of the fraction (Fig. 2A). This antibody depletion in the FT was reflected in the MIB-captured fraction, where antibodies constituted ∼90% of the total protein abundance. The remaining 10.4% of the MIB fraction contained low-abundance contaminants, e.g., hemopexin, albumin, and serpin A1, which were approximately 10 times less abundant than the antibodies (File S2). Of note, the most abundant contaminants align well with the most abundant proteins in the pooled serum control. Importantly, the MIB pull-down yielded a near-complete recovery of all main antibody isotypes in the eluate, capturing 93.2% of IgM and more than 99% of all IgG and IgA isotypes from human serum (Fig. 2B). These results suggest that MIB universally binds all antibodies with high affinity and does not seem to discriminate between antibody isotypes and subclasses.

**Figure 2.**
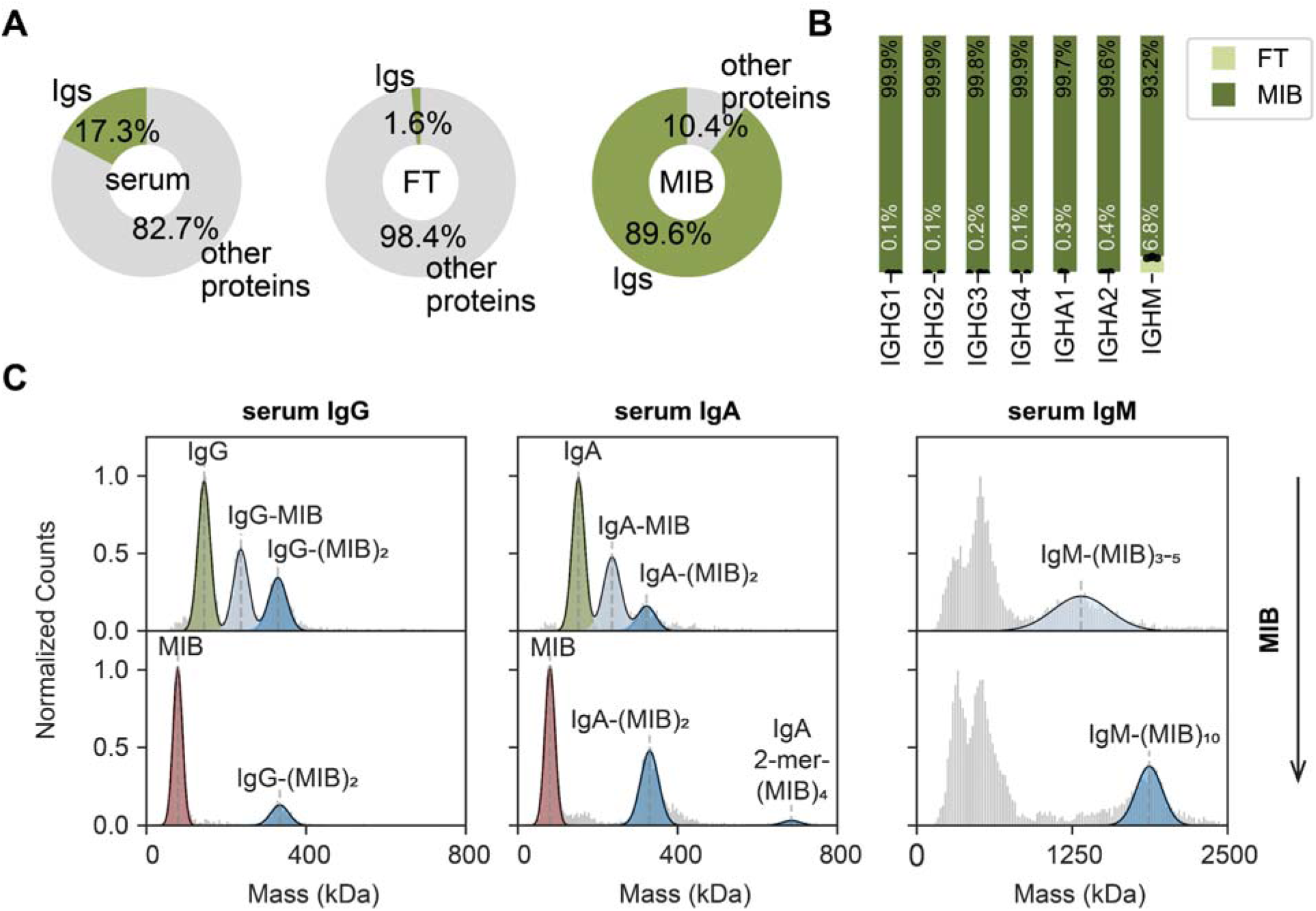
MIB universally and selectively binds human immunoglobulins. **A** DIA LC-MS/MS based quantitative proteomics of antibody abundance in the MIB affinity capture experiments from pooled donor human serum. The pie charts depict relative protein abundances in the serum control, MIB-unbound flow-through (FT), and MIB captured fraction, from left to right. The data suggest that MIB effectively depletes all antibodies from the serum sample. Conversely, antibody enrichment is observed in the MIB-captured fraction. Percentages in the pie chart represent means of n=4 replicates. **B** Distribution of IgG, IgA, and IgM (sub)classes in MIB-unbound FT (light green), and the MIB-captured fraction (dark green) from **A** as quantified by DIA LC-MS/MS based quantitative proteomics. The results correspond to means of n=4 analyses; individual replicates are indicated by black dots. **C** Mass analysis of MIB binding to human serum-derived polyclonal antibodies by mass photometry. The histograms depict normalized distributions of single-particle landing events by mass (kDa) for polyclonal, serum-derived IgG, IgA, and IgM with MIB. MIB (red) can bind human antibodies (green) universally, forming antibody–MIB complexes in various stoichiometries (blue). The unfitted densities in the IgM panel correspond to common, previously reported, contaminants^8^ that are co-isolated from serum and do not interact with MIB. The IgG sample was purified from serum from a single healthy donor; IgM and IgA were purified from pooled sera from healthy donors (Amsbio). The results were acquired with the following antibody–MIB ratios: top row (1:1, 1:2, and 1:5), bottom row (1:5, 1:8, and 1:20). Measured and expected masses of annotated species are included in Table S1.

To further validate this observation and directly monitor MIB binding to serum-derived polyclonal antibodies, we probed the interactions by single-particle mass photometry (MP).^30^ The MP results confirmed that MIB is an 85 kDa soluble protein that binds polyclonal IgG, IgA, and IgM (Figs. 2C, S1, Tab. S1–2). While for IgG and monomeric IgA we observed 1:1 and 1:2 antibody:MIB complexes, J-chain coupled IgA dimers also formed 1:4 complexes. The pentameric IgM bound up to 10 MIB molecules. Notably, all serum-derived polyclonal IgM samples contain low-mass protein contaminants (Fig. S1–2), which were previously observed^8^ and did not interact with MIB, highlighting the specificity of MIB’s interaction with immunoglobulins. These results suggest that all Fab domains within an antibody molecule can be simultaneously occupied by MIB, independent of the antibody isotype (Fig. 2C), resulting in two MIB molecules bound per antibody protomer. This general observation could be further validated for polyclonal human colostrum-derived SIgA containing pIgR, and several other polyclonal IgA and IgM samples (Fig. S2, Tab. S2).

To assess whether the structural complexity or sequence variability of antibodies influences MIB binding, we further analyzed MIB complexes with several recombinant monoclonal antibodies. Starting with the potential influence of the oligomeric state, we compared the interaction of MIB with three isotypic variants of a monoclonal recombinant anti-WTA 4497: IgG, dimeric J-chain coupled IgA, and pentameric J-chain coupled IgM. The oligomeric species were obtained through isotype conversion of the anti-WTA 4499 monoclonal antibody, allowing us to assess the effects of structural differences among isotypes exclusively. We performed MIB titrations while maintaining a constant antibody protomer concentration across all isotypes and analyzed and quantified the resulting complexes by MP (Fig. 3). Binding followed a similar pattern across all three antibody isotypes. We observed that half of the Fabs were occupied at approximately 1.5 MIBs per protomer and a plateau at 2 MIBs per protomer in the presence of a slight MIB excess (∼2.5 MIBs per protomer). Thus, MIB can effectively saturate the antibodies regardless of the oligomeric state, as visualized in Fig. 3B and in Fig. 2C for polyclonal samples.

**Figure 3.**
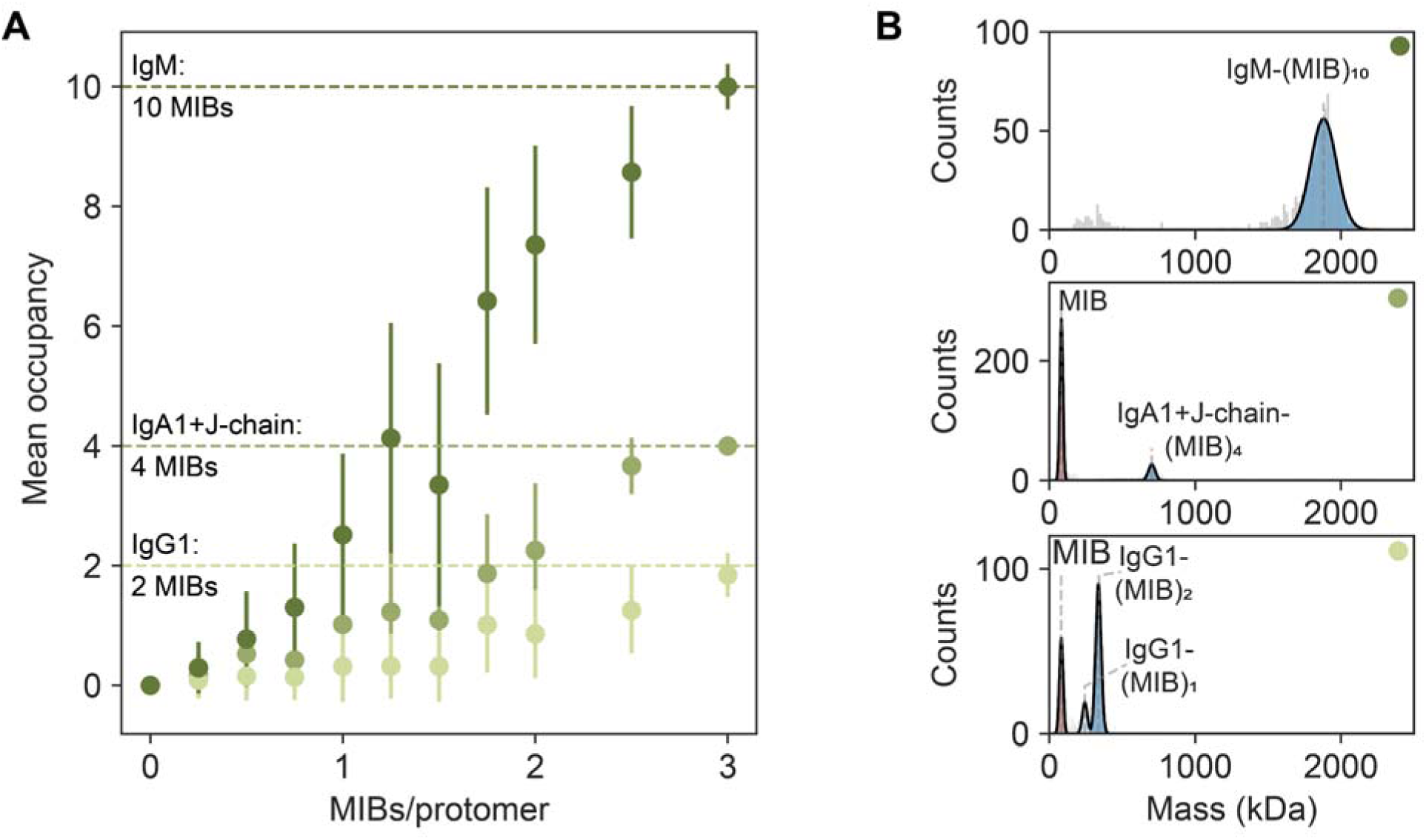
Two MIB molecules bind each antibody protomer. MIB interaction with monoclonal anti-WTA 4497 antibodies examined by mass photometry: IgG1 (light green, 1 protomer), IgA1+J-chain (medium green, 2 protomers), and IgM+J-chain (dark green, 5 protomers). **A** Mean occupancy of each immunoglobulin acquired at various MIB/protomer ratios in range of 0**–**3. The results suggest that each of the immunoglobulins reaches full occupation of 2 MIBs/protomer, regardless of the polymeric state. The lines correspond to the standard deviation of the mean occupancy for IgG and IgA+J and relative sigma in case of IgM. **B** Representative mass photometry histograms for IgM, IgA1, and IgG1 at 3 MIBs/protomer used for the quantification depicted in panel **A**. The histograms show full occupation of each antibody by MIB (red) forming fully saturated complexes with 2 MIBs/protomer (blue). Measured and expected masses of annotated species are included in table S1.

Next, we systematically assessed whether sequence variations within a single isotype or across isotypes cause differences in MIB binding (Figs. S3–S4, Tab. S2). We started with a panel of IgG1 antibodies targeting different epitopes, carrying either κ or λ light chains, and Fab glycosylation. We found that none of these traits affected the interaction with MIB. The analysis of monomeric IgG2–4 and IgA1–2 yielded similar results, suggesting that neither the subclass nor the isotype of monomeric antibodies affects MIB binding (Fig. S3). Consistent with the aforementioned results (Fig. 3), we did not observe an influence of the oligomeric state, J-chain, or CD5L presence on the tight antibody–MIB interaction (Fig. S4, Tab. S2). These MP observations were further confirmed by differential scanning fluorimetry (DSF), as presented in Fig. S5. Our DSF results exhibited similar effects on the stabilization and melting temperatures across all antibody–MIB complexes, regardless of isotype or oligomeric state. In summary, MIB binding appears unaffected by sequence or structural differences and reaches an occupancy of 2 MIBs per protomer across all examined antibodies, further positioning MIB as a universal binder of human antibodies.

To further investigate the structural basis of MIB recognition that allows its broad specificity across antibody isotypes, we employed cross-linking mass spectrometry (XL-MS) and high-speed atomic force microscopy (HS-AFM). We performed these analyses with recombinant monomeric IgG1 and recombinant pentameric J-chain coupled IgM to cover the breadth of antibody structures (Figs. 1, 4). HS-AFM analysis of the antibody–MIB complexes revealed high occupancy with mostly two MIBs binding per antibody protomer (Fig. 4A–B, Tab. S3), consistent with the MP data. While free MIB formed approximately 20 nm globular particles with a flexible domain and a height of 4.2 nm, free antibodies were detected as particles composed of Y-shaped protomers with Fab heights of 4.0–4.4 nm (Fig. S6, Tab. S3), in agreement with previous observations.^15,31^ The antibody–MIB complexes revealed MIB binding to the Fab regions, as evidenced by an increase in Fab height to 7.1–7.2 nm. This is ∼1.3 nm less than the cumulative height of MIB with Fab, indicating substantial structural changes upon antibody–MIB interaction.

**Figure 4.**
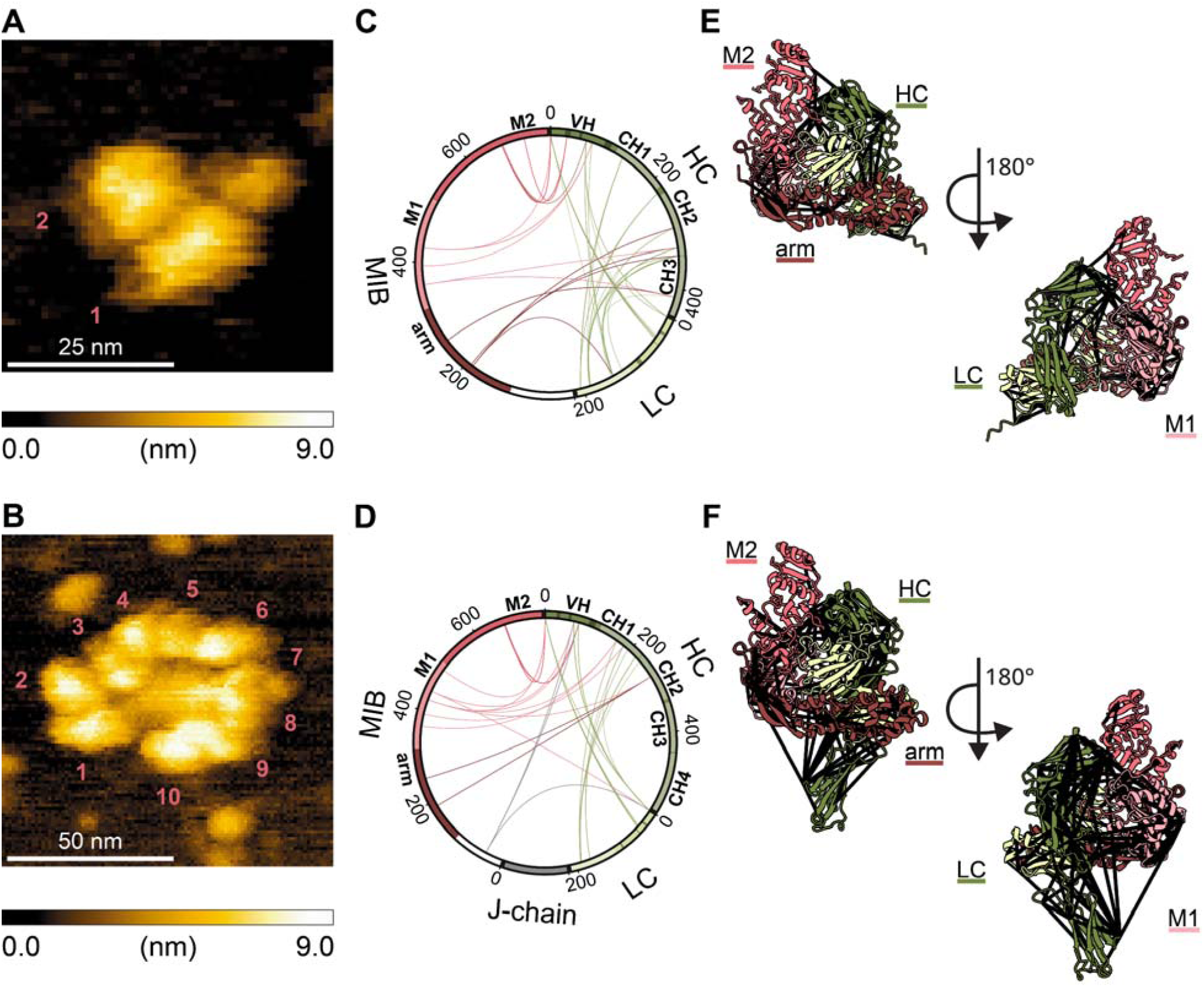
MIB binds IgG and IgM in a similar manner. **A,B** High-speed atomic force microscopy of anti-WTA 4497: **A** IgG1 with 2 MIBs and **B** IgM with 10 MIBs immobilized on mica surface. The images reveal the involvement of antibody Fab regions in MIB binding. Each individual MIB–Fab interaction binding is labeled by the depicted numbers. **C, D** Cross-linking MS of **C** trastuzumab IgG1–MIB and **D** CoV07 IgM–MIB complexes. Each line in the circos plots corresponds to a PhoX inter-protein cross-link detected in all replicates with at least 7 cumulative CSMs. The intra-antibody cross-links connecting the heavy chain (HC, dark green) and light chain (LC, light green) are shown in green, while the MIB-antibody cross-links are colored based on MIB-domain involved in the interaction: arm (111–319, dark red), M1 (320–510, light pink), M2 (511–749, salmon). **E, F** Visualization of the cross-linking MS data on AlphaFold3-based structural models of MIB interaction with Fab regions of **E** IgG1 (trastuzumab) and **D** IgM (CoV07). The colors match **C, D**, all cross-links are shown as black lines. The cross-links reveal a similar basis for MIB interaction with IgG and IgM. While the MIB M2 domain engages with the VH of the antibody, the MIB arm interacts predominantly with the CH2 domains.

To obtain more localized insights into antibody–MIB binding, we performed XL-MS (Figs. 4C–D, S7, File S3). XL-MS uses covalent reagents to map residues that are spatially close in protein structures or directly involved in protein complex formation.^32,33^ The XL-MS data acquired with the PhoX chemical cross-linker showed that MIB binds trastuzumab IgG1 (Fig. 4C, E) and CoV07 IgM (Fig. 4D, F) in a comparable manner, despite the differences in antibody sequences and structures (Fig. 1). Importantly, the intra-antibody cross-links (green) corroborate interactions between the constant domains of the heavy and light chains, as well as between the variable domains of both chains. The antibody–MIB interaction was predominantly captured by restraints involving the heavy chain. For both antibodies, the C-terminal MIB M2 domain engaged with framework region 3 of the antibody heavy variable domain. Despite some differences in MIB M1 domain intra-links, we detected analogous cross-links to the first framework region and to the constant domains of the heavy chain. Lastly, cross-links were observed between the N-terminal MIB arm and the CH2 domain of both IgG1 and IgM, indicating that this domain localizes closer to the Fc portion. To visualize the XL-MS data and MIB binding to IgG and IgM, we generated structural models of the Fab–MIB interactions (Fig. 4E–F), which align well with earlier reported cryo-EM structure of MIB with polyclonal goat IgG (Fig. S8).^28^ The models highlight interaction interfaces captured by XL-MS. Specifically, we observed contacts between the MIB M2 domain and the variable region of the heavy chain, and between the MIB arm and the constant region of the heavy chain. The model’s compatibility with the XL-MS results is also consistent with the restraint distance distributions for both IgG and IgM (Fig. S8).

### MIB disrupts antibody Fab structures without affecting Fc effector functions

The HS-AFM movies of free and MIB-bound IgM (Mov. S1–2) exposed another aspect of antibody–MIB interactions. While the Fab arms of the free IgM readily fluctuated over the course of the measurement, the MIB-bound Fabs appeared substantially more rigid. To quantify these effects, we tracked trajectories of free and MIB-bound Fabs over the course of the measurement (Fig. 5A). Already, the particle trajectories showed more restricted movement of MIB-bound Fabs when compared to Fabs of free IgM. To directly compare and quantify the impact of MIB binding, we performed mean square displacement (MSD) analysis (Fig. 5B, Tab. S4). The results show that both free (green) and bound (blue) Fabs follow the confined diffusion model. Yet, the radius of circular confinement was approximately 2.6 times smaller for the MIB-bound complex, indicating restricted flexibility. To confirm that these results are also applicable to other antibody isotypes, we investigated IgG1-RGY, a recombinant IgG1 that readily hexamerizes in solution (Mov. S3, Fig. 5C–D, Tab. S4).^11^ The results revealed that, also for the IgG1-RGY hexamer, Fab mobility dramatically reduces upon MIB binding, suggesting that MIB binding generally reduces Fab flexibility regardless of the antibody isotype.

**Figure 5.**
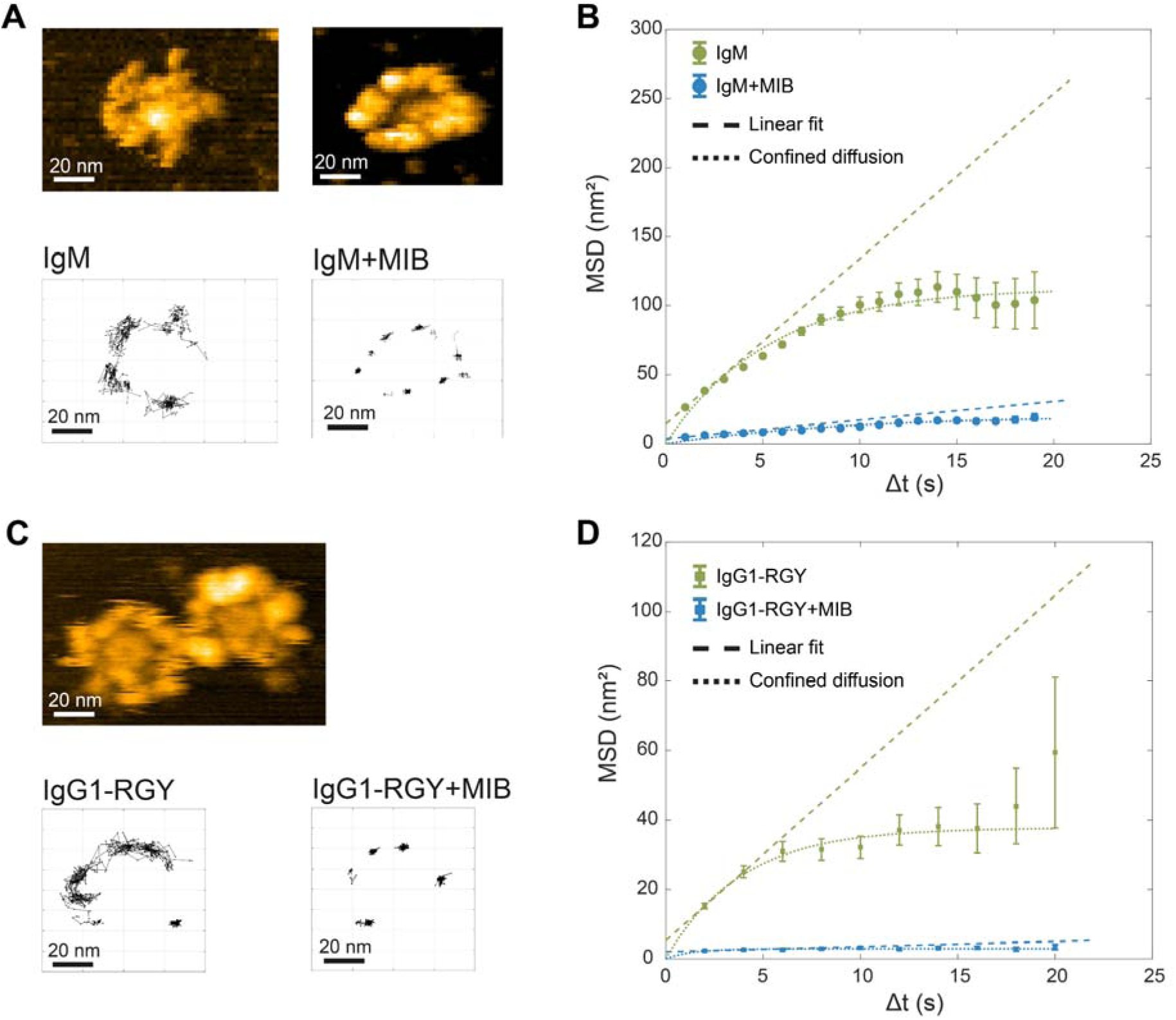
MIB binding confines Fab flexibility. Time-resolved high-speed atomic force microscopy of **A, B** 4497 IgM and **C, D** CD52 IgG1-RGY hexamers, either free or in complex with MIB. **A, C** Representative images from high-speed atomic force microscopy movies and corresponding Fab domain trajectories covered for free (left) or MIB-bound (right) antibodies. **B, D** Mean square displacement (MSD) analysis of Fab domain trajectories. The MSD(Δt) is described by a confined diffusion model (dotted line). A linear fit to the first two MSD data points characteristic of free diffusion (dashed line) is included for comparison. The results suggest that MIB-bound Fab domains (blue) are less flexible than those of free antibodies (green) for both **A, B** IgM and **C, D** IgG1-RGY. Results of the linear and confined diffusion fit in **B**, **D** are included in Tab S4.

The HS-AFM results revealed that MIB binding affects Fab structure and flexibility. In contrast, we observed that MIB does not impair Fc-mediated hexamerization of IgG1, suggesting that MIB’s impact on antibody structure may be confined to the Fab domain. To further probe this effect, we next aimed to characterize MIB’s influence on the formation of antibody complexes. As anticipated, MIB efficiently disrupted antibody–antigen complexes of IgG1 (Fig. 6A),^28^ as well as those of IgM with mono- and polyvalent antigens (Fig. 6B, S9). Moreover, when antibodies were pre-incubated with MIB, antibody–antigen complex formation was prevented (Fig. S9). In all cases, no simultaneous interaction of a single Fab arm with both MIB and antigen was observed.

**Figure 6.**
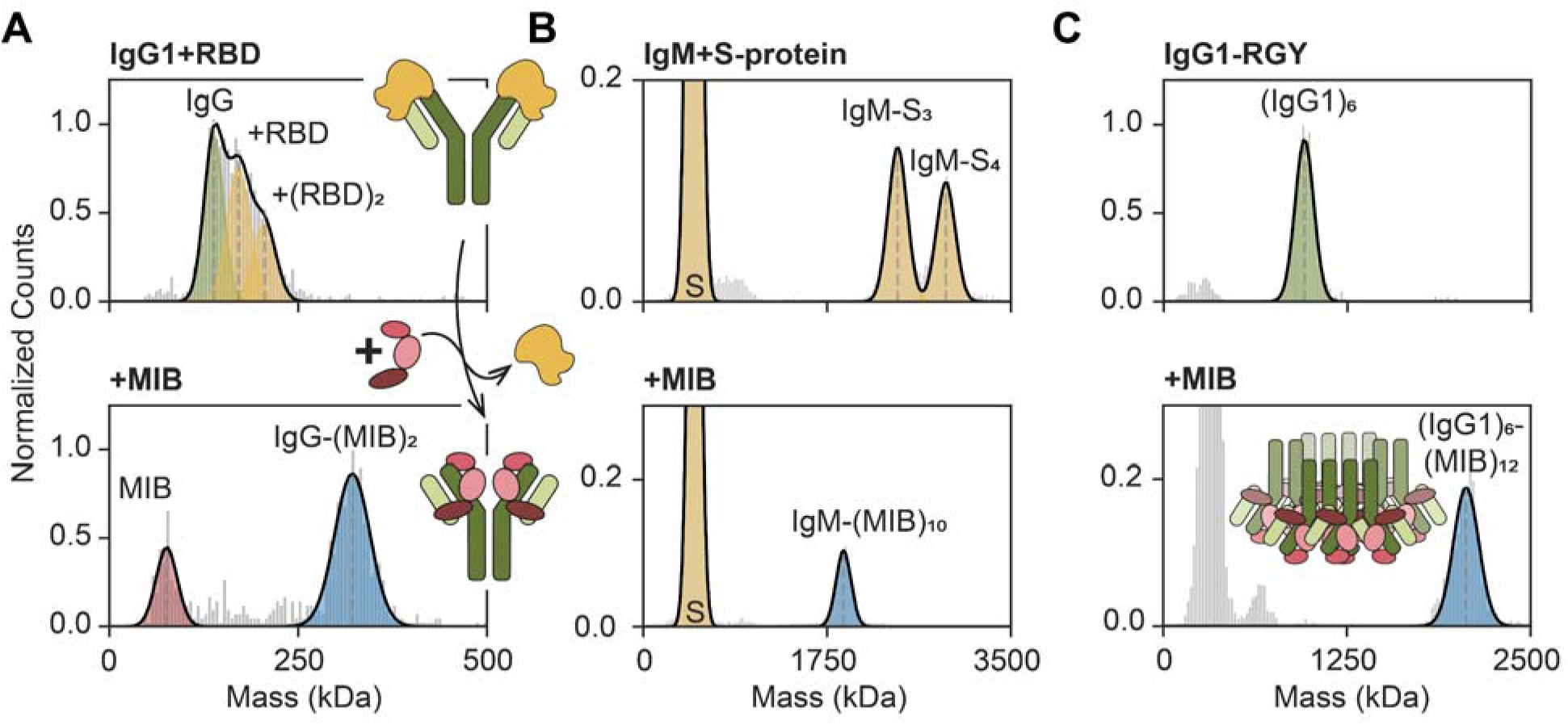
MIB disrupts antibody–antigen immune complexes and can interact with IgG hexamers. Mass photometry histograms displaying the interactions of antibody complexes with MIB. **A** Top panel shows anti-S-protein RBD IgG1 CoV07 (green) bound to RBD antigen (orange). The bottom panel shows the same sample incubated with MIB, highlighting MIB-mediated disruption of the IgG1–RBD complex through formation of the IgG1–MIB complex. **B** Top panel shows anti-S-protein RBD IgM CoV07 bound to S-protein antigen (orange). The bottom panel highlights the disruption of the IgM–S-protein complex by MIB addition through the formation of the IgM–MIB complex. **C** Campath anti-CD52 IgG1-RGY hexamers (green, top panel) and 18-component assembly of MIB-bound IgG1 hexamer (blue, bottom panel). The MP analysis suggests that while MIB disrupts antibody–antigen complexes, it does not interfere with Fc-mediated protein interactions. Unannotated low MW intensities in **C** correspond to overlapping signals of MIB and IgG1 protomers, which are below the detection limit. The data were acquired with the following protein complex component abundances: antibody (1 mol. eq.), **A** RBD (5 mol. eq), MIB (5 mol. eq.), **B** S-protein (10 mol. eq.), MIB (20 mol. eq.), **C** MIB (20 mol. eq.). Masses and expected masses of annotated species are included in table S1.

To examine the effects of MIB on Fc-mediated interactions, we used hexameric IgG1-RGY constructs and characterized their interactions with excess MIB (Figs. 6C, S10). The MP analysis revealed the formation of ∼1.9 MDa complexes containing six copies of IgG1 (6 × 148 kDa) and twelve copies of MIB (12 × 85 kDa), following the general trend of 2 MIBs/protomer. It was previously reported that IgG1-RGY hexamers can efficiently bind to complement component 1q (C1q) in solution.^11,34^ C1q is a large 460 kDa protein complex composed of six copies of α-, β-, and γ-chains, which serves as the first component of the classical complement pathway.^35^ Thus, we next assessed whether the MIB-bound IgG1 hexamers could still bind C1q. The data revealed that C1q still interacts with MIB-decorated IgG1 hexamers, forming a ∼2.3 MDa assembly (Figs. 7A, S11). Moreover, using soluble CD38 antigen (30 kDa), we observed that MIB can also disrupt antibody–antigen interactions in the presence of C1q (Fig. 7A–C). This was evidenced by a ∼0.66 MDa mass shift in the mass histograms corresponding to the displacement of 12 CD38 antigen molecules by 12 MIB molecules. A similar mass shift was observed for the IgG1-RGY complex in the absence of C1q. Taken together, these results indicate that MIB selectively displaces antigens even when incorporated into large immune complexes, while retaining Fc-mediated interactions and thus likely preserving the ability to activate complement.

**Figure 7.**
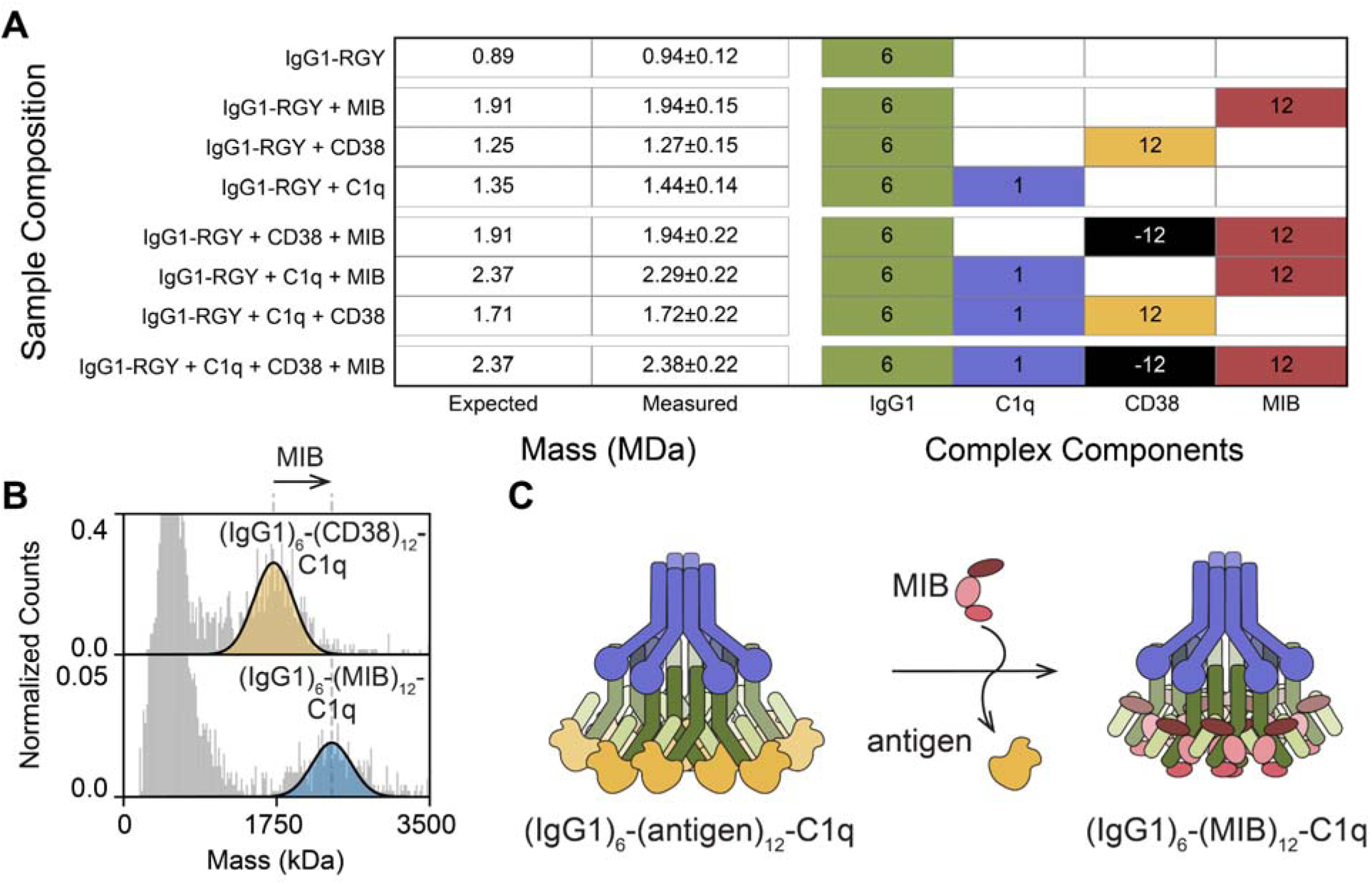
MIB displaces antigens in oligomeric immune complexes. **A** Interaction of anti-CD38 005-IgG1-RGY hexamers with C1q, soluble CD38 antigen, and MIB mapped by mass photometry. The table shows a summary of mass photometry experiments, including expected and measured mass (MDa) of detected high mass species. The components forming respective high mass complexes: IgG1 (green), C1q (purple), CD38 (orange) and MIB (red) are included with copy numbers. Antigen displacement, evidenced by a ∼0.66 MDa mass shift, as observed for samples IgG1-RGY + CD38 + MIB and IgG1-RGY + C1q + CD38 + MIB, is highlighted in black. **B** Normalized mass histograms of IgG1-RGY + C1q+ CD38 (top), and IgG1-RGY + C1q + CD38 + MIB evidencing (bottom). The data shows antigen displacement from (IgG)_6_-(CD38)_12_-C1q by MIB and formation of the (IgG)_6_-(MIB)_12_-C1q complex (∼2.3 (∼2.3 MDa) as illustrated in cartoon in **C**. The data were acquired with following molar ratios of each complex component: IgG1-RGY (1 mol. eq.), C1q (4 mol. eq.), MIB (20 mol. eg.), CD38 (20 mol. eq.).

## Discussion

Here, we characterized MIB as a pan-antibody-binding protein, demonstrating universal high affinity for all main human antibody isotypes. Using MIB, we efficiently co-depleted IgG, IgA, and IgM from human serum, with over 99% capture efficiency for IgG and IgA and 93% for IgM. Near-complete antibody capture, along with low levels of co-purified contaminants, underscores MIB’s specificity for antibodies and its unbiased behavior across isotypes. This was mirrored in the characterization of MIB interactions with a diverse panel of recombinant monoclonal antibodies and endogenous polyclonal antibodies. MIB efficiently bound all studied antibodies regardless of their oligomeric state, sequence variability, light chain type, Fab glycosylation, or incorporation of J-chain, CD5L, or pIgR. Moreover, MIB consistently saturated antibodies with two MIB molecules per protomer regardless of their structural features, as confirmed by both MP and HS-AFM. This suggests that MIB binding is guided by highly conserved structural features of antibodies.

Interactions of IgG and IgM with MIB were further structurally probed by XL-MS. Cross-links marked interactions with MIB that were consistent between antibody isotypes. The observed interactions of MIB with the first and third framework regions of the heavy variable domains, as well as with the constant domains of the heavy chain, support the hypothesis that MIB interacts with highly conserved regions. Our cross-links align well and extend beyond the available cryo-EM data on a goat IgG Fab domain, which shows that MIB encircles the light variable domain.^28^ Notably, cross-links were observed between the N-terminal MIB arm and the CH2 domain of both IgG1 and IgM. Simultaneous interactions with the Fab and CH2 domains may lock the Fab domain in place, which could induce the dramatic reduction of Fab structural flexibility observed for IgM and hexameric IgG1-RGY by HS-AFM. Increased rigidity of the Fab region may be vital for MIP recruitment or to position the Fab for proteolytic cleavage, since MIP does not interact without MIB.^20,28^ Moreover, the observed structural rigidity of MIB-bound Fab domains could aid in disrupting antibody–antigen complexes, a process previously attributed solely to conformational shifts in the complementarity-determining regions.^28^

Interestingly, the functional effects of MIB interactions appear to be localized to the Fab domain, as the effector functions of the Fc domain remain largely intact. MIB does not affect the hexameric IgG1-RGY composition that is established through interaction of the Fc chains.^36^ Moreover, C1q can still interact with MIB-bound IgG1 hexamers, forming an (IgG)_6_-(MIB)_12_-C1q immune complex, and can even disrupt antibody–antigen interactions in the presence of C1q. Formation of these large immune complexes devoid of antigen could hint at a mechanism underlying the chronic inflammatory nature of mycoplasma infections. To better understand this, additional functional studies are required to determine whether MIB interferes with complement pathway activation.

To conclude, MIB is an antibody–binding protein that universally recognizes all main human antibody isotypes and subclasses with high affinity. Interestingly, MIB shows a different binding profile compared to other bacterial antibody binders, such as S-protein A and protein G, which preferentially bind the IgG isotype.^37^ This difference can be attributed to MIB targeting conserved regions in the Fab domain rather than the Fc domain, also allowing MIB to incorporate into large immune complexes. Since protein A and G are widely used in biotechnological applications, we propose that MIB could be a valuable addition to the current technological toolset. The pan-antibody-binding profile and high-affinity position MIB as an effective tool for pan-antibody profiling or for efficient depletion of antibodies from serum.

## Materials and Methods

### Materials, Chemicals, and Protein Samples

Unless otherwise stated, all experiments were conducted using Milli-Q H_2_O generated by the IQ 7003 system (Merck, Germany). PhoX cross-linker was synthesized as described before.^38^ Dimethyl sulfoxide (DMSO), formic acid (FA), 4-ethylmorpholine, and trifluoroacetic acid (TFA) were acquired from Thermo Scientific, USA. Trypsin was obtained from Promega, USA, and LysC from Wako, Germany. Chloroacetamide (CAA), cell culture phosphate-buffered saline (PBS), sodium deoxycholate (SDC), Trizma Pre-set crystals (pH 8.5), Tris(2-carboxyethyl)phosphine (TCEP), and urea were acquired from Merck, Germany. LC-MS solvents A (0.1% v/v FA in H_2_O) and B (80% acetonitrile (ACN) v/v, 0.1% FA v/v in H_2_O) were acquired from Biosolve Chimie, France.

MIB protein from *M. hominis* was recombinantly expressed in *E. coli* and provided by Genovis AB, Sweden. Cetuximab IgG1 was acquired from Evidentic, Germany. F95 IgG1 was recombinantly expressed as described before.^39^ Trastuzumab was provided by Roche, Germany. 4497-IgG1, -IgG2, -IgG3, -IgG4, -IgA1+J-chain, IgM (± J-chain and CD5L) were recombinantly expressed, as described before.^8,40^ Anti-StrepTagII IgM+J-chain was recombinantly expressed, as described before.^41^ 2F8-IgG1, -IgG1-RGY, -IgG2, -IgG3, -IgG4, - IgA1; 5d5-IgA2; 005-IgG1-RGY; Anti-CD52-IgG1-RGY; soluble epidermal growth factor receptor (EGFR); and soluble CD38 were kindly provided by Genmab and recombinantly expressed as described before.^34,42,43^ Anti-S-protein CoV07 IgM and IgG1 were obtained from AstraZeneca, UK. 2P-stabilized spike (S) protein of the Wuhan strain of SARS-CoV-2 and respective receptor-binding domain (RBD) were recombinantly expressed as described before.^44^ Hepatitis C soluble envelope protein sE1E2 with StrepTagII was recombinantly expressed as described before.^45^ Human C1q was acquired from Complement Technology, USA.

A polyclonal IgG sample was purified from a single healthy donor serum using the CaptureSelect™ FcXL Affinity Matrix (Thermo Scientific, USA) as described previously.^46,47^ Polyclonal IgA samples were purified from pooled healthy donor serum (Amsbio, UK) and single healthy donor plasma SER-PLE (Zen-bio, USA) using CaptureSelect™ IgA-XL Affinity Matrix (Thermo Scientific, USA), as described before.^48^ Human colostrum-purified SIgA was acquired from Sigma-Aldrich. Polyclonal IgM was purified from healthy donor serum (Amsbio, UK) using the CaptureSelect™ IgM-XL Affinity Matrix (Thermo Scientific, USA), as previously described.^49^ Additionally, human plasma-purified IgM samples were acquired from Prospec and Sigma-Aldrich.

The presented data were partially post-processed using the following Python libraries: matplotlib, NumPy, Pandas, pyCirclize, SciPy, and seaborn.^50–53^

### Affinity pull-down using immobilized MIB

1 mg of recombinant MIB was immobilized onto 0.4 mL of Pierce NHS-Activated Agarose (Thermo Scientific, USA) following the vendor-provided protocol. The slurry with immobilized MIB was divided into four Pierce Micro-Spin Columns (Thermo Scientific, USA), 100 μL of MIB slurry/column. These MIB columns were subsequently used to perform affinity pull-down from pooled donor healthy human serum (Amsbio, UK) to identify MIB interactors. The experiment was performed in n=4 replicates.

First, the columns were washed with 200 μL PBS (3×, 1000 × g, 1 min) and incubated for 2 h at room temperature (RT) at 800 rpm with 10 μL of pooled donor human serum diluted with 190 μL PBS. The flow-through (FT) was collected, and columns were washed with 200 μL PBS (3×, 1000 × g, 1 min). Afterward, 200 μL of 0.1 M glycine, pH 2.7, was added, and after 10 min incubation at RT, 800 rpm, the MIB-captured fraction was collected. The eluted sample was neutralized with 10 μL of 1 M Tris, pH 8, and all samples, including an untreated serum control, were subjected to bottom-up proteomics analysis.

### Bottom-up proteomics

The protein concentration in each fraction was determined by the Bradford protein assay using Protein Assay Dye Reagent (Bio-Rad, USA), with bovine serum albumin (BSA) as the standard, on a Multiskan GO plate reader (Thermo Scientific, USA). The assay was performed according to the vendor-provided protocol, and 1 μg of each sample was digested. The samples were denatured, reduced, and alkylated in SDC buffer for 30 min at RT (final concentration: 0.1 M Tris, pH 8.5; 40 mM TCEP; 100 mM CAA; 1% SDC (w/v)). Subsequently, the samples were diluted 10× with 50 mM ammonium bicarbonate, pH 8.5, and digested with trypsin (1:50 w/w) and LysC (1:75 w/w) for 16 h at 37 °C. The reaction was stopped by adding TFA (0.5% v/v, final concentration), and the SDC precipitate was removed by centrifugation (20 000 × g, 10 min).

Finally, the samples were desalted on an Oasis HLB μElution Plate (Waters Corp, USA) using a vendor-provided protocol. The solvent was evaporated in a vacuum centrifuge and stored at -20 °C. The dry peptides were resuspended in 2% FA (v/v) prior to the LC-MS/MS analysis.

The bottom-up proteomics analysis was performed in data-independent acquisition (DIA) mode on an Orbitrap Exploris 480 and an Ultimate 3000 UHPLC (both Thermo Scientific, USA). The peptides were analyzed in positive ion mode using an 85 min method, starting with a 1 min trapping step at 30 μL/min, with solvent A (0.1% FA v/v in H_2_O), on an Acclaim Pepmap 100 C18 column (5 mm × 0.3 mm, 5 μm, Thermo Scientific, USA). The peptides were separated on an in-house packed analytical column (Reprosil 2.4 μm, 75 μm × 50 cm) at a flow rate of 0.3 μl/min. The following analytic gradient was used for solvent B (80% ACN v/v; 0.1% FA v/v in H_2_O): 1 min - 9% B; 2 min - 13% B; 67 min - 44% B; 70–75 min - 99% B; 75–85 min - 9% B. The MS1 scans were acquired over a 375–1600 m/z range at a resolution of 60 000. The DIA MS2 scans were acquired at a 3s cycle time with 30 windows over the 400–1000 m/z range, 20 m/z windows, and 1 m/z overlap. Each MS2 scan was acquired with a 30,000 resolution, 28% normalized HCD, standard AGC, and automatic injection time.

The data were processed in library-free mode with DIA-NN (2.0).^54^ The peptide length was set to 7–30 amino acids, the precursor m/z range was 300–1800, and the charge range was 1–4. One missed cleavage was allowed, carbamidomethyl was set as a fixed modification, and methionine oxidation was allowed as a variable modification. The reviewed human proteome with isoforms (UniProt, 09/2023), including common contaminants, was used as a database. The results were filtered for 1% FDR and contaminants. Only proteins with at least 2 peptides were included, and LFQ was recalculated only using peptides without oxidation as described before (File S1).^55^ The immunoglobulins in each sample were quantified based on median LFQ values. The antibody heavy and light chain genes, as well as J-chain, CD5L, and pIgR, were included as immunoglobulins (File S2), and the immunoglobulin sum median intensity was compared with the total median intensity in each sample.

### Mass photometry

The MP data were acquired on the OneMP or the SamuxMP (both Refeyn Ltd, UK), depending on the expected molecular weight (MW) of the analytes. Particularly, all IgM and IgG1-RGY complexes were analyzed on SamuxMP, since the expected MW exceeded 1 MDa. Analyses of IgG1–4 and IgA1–2 and respective complexes were performed on the OneMP. All analyses were performed in PBS, pH 7.4.

The antibody–MIB titration series (Fig. 3) was executed using anti-WTA 4497: IgG1, IgA1+J-chain, and IgM to achieve a final 32 nM antibody protomer concentration (i.e., 32 nM IgG1, 16 nM IgA1+J-chain, and 6.4 nM IgM). MIB was added at concentrations ranging from 0.25 mol. eq. (8 nM, final concentration) to 3 mol. eq. (96 nM, final concentration). The data presented in Fig. 1, 6–7, S2, S4, S9–10, and Tab. S1–2 were generated similarly, with depicted MIB:antibody ratios. The MIB amounts were selected based on Fig. 3 to achieve full antibody occupation. Unless otherwise stated, all complexes were incubated at RT for at least 30 min prior to measurement.

The MP analysis was performed as previously described.^43,56^ Briefly, 12 μL of buffer was pipetted on an in-house carrier slide and used for the automatic focusing procedure. Subsequently, 1–3 μL of the sample was added to the buffer to achieve the optimal concentration for single-particle measurement. Raw MP movies were recorded for 60 s at 100 frames per second and mass-calibrated in DiscoverMP (Refeyn Ltd, UK). The OneMP calibrant mixture consisted of IgG-halfbody (73 kDa), IgG (149 kDa), apoferritin (513 kDa), and IgM (1006 kDa). SamuxMP calibration was performed using bovine thyroglobulin oligomers (335 kDa, 670 kDa, 1005 kDa, 1340 kDa). The calibrated events were exported and further analyzed.^43,56^ The resulting mass distributions were plotted with a bin size of 20 kDa for SamuxMP and 5 kDa for the OneMP. The masses and species abundances were determined using a multi-Gaussian fit, based on selected starting parameters. The mean MIB occupancies of samples with resolved single MIB binding events (IgG, IgA) were calculated as weighted averages of species abundances with standard deviation (SD). The mean IgM occupancies were determined from the mass difference of the MIB-bound complex with the free IgM control divided by the MIB mass. The relative sigma corresponds to the sigma (SD) of the Gaussian distribution relative to the sigma of the free IgM control.

### Cross-linking mass spectrometry

XL-MS was performed in experimental triplicate using PhoX cross-linker.^38^ First, the optimal cross-linker concentration was determined by reducing SDS-PAGE (Fig. S11), and 1.5 mM PhoX was selected for the experiment. MIB was mixed with recombinant monoclonal antibodies trastuzumab IgG1 or CoV07 IgM+J-chain to achieve a ratio of 2 MIB molecules per antibody protomer. This mixture (0.1 μg/μL in PBS, pH 7.4) was incubated for 20 min at RT prior to the XL reaction. The reaction was initiated by adding PhoX dissolved in DMSO, resulting in a final DMSO concentration of 1.5% (v/v) in the sample. The reaction was terminated after 1h of incubation at RT by the addition of 25 mM Tris, pH 8 (final concentration), and incubated for an additional 30 min at RT. The resulting samples were first denatured with 6 M urea (final concentration), then 3× diluted with 50 mM 4-ethylmorpholine, pH 8.5; 40 mM TCEP; 100 mM CAA, and incubated for 30 min at RT. The digestion was performed with trypsin (1:50 w/w) and LysC (1:75 w/w) at 37 °C for 16 h. The digestion was terminated with TFA (0.5% v/v, final concentration), and samples were desalted on an Oasis HLB μElution Plate (Waters Corp, USA) using a vendor-provided protocol and fully evaporated in a vacuum centrifuge. The peptides were deglycosylated with Protein Deglycosylation Mix II (New England Biolabs, USA) using the vendor-provided non-denaturing protocol and subsequently subjected to phosphopeptide enrichment using Bravo AssayMAP liquid handling robot (Agilent, USA) with AssayMAP 5 µL Fe(III)-NTA cartridges (Agilent, USA) using vendor-provided protocol.^38^ The phospho-enriched fraction was fully evaporated and dissolved in 2% FA (v/v) prior to the analysis.

The LC-MS/MS experiment was performed in data-dependent acquisition (DDA) mode on an Orbitrap Exploris 480 with an Ultimate 3000 UHPLC (both Thermo Scientific, USA). Peptides were analyzed in positive mode using a 70 min method starting with trapping 1 min, 30 μL/min, solvent A (0.1% FA v/v in H_2_O) on an Acclaim Pepmap 100 C18 (5 mm × 0.3 mm, 5 μm, Thermo Scientific, USA). The peptides were separated on an in-house packed analytical column (Reprosil 2.4 μm, 75 μm × 50 cm) at a flow rate of 0.3 μl/min. The following analytic gradient was used for solvent B (80% ACN v/v; 0.1% FA v/v in H_2_O) percentages: 1 min - 9% B; 2 min -13% B; 57 min - 41% B; 58–62 min - 99% B; 63–70 min - 9% B. The MS1 scans were acquired in a 375–2200 m/z range with a 60 000 resolution. The DDA MS2 scans were acquired for charge states 3–8 in a 2 s cycle time with a 15 s exclusion after single precursor selection using the following parameters: 120 m/z first mass; 1.4 m/z isolation window; 30 000 resolution; stepped 20, 28, 36% normalized HCD; standard AGC; and 25 ms maximum injection time.

The resulting files were processed with pLink 3.0.16^57^ using a database containing MIB, respective antibody sequences, as well as common contaminants provided by pLink 3.0.16. The following settings were used: protease – trypsin (with 3 allowed missed cleavages); peptide mass – 600–6000 Da; peptide length – 6–60; cross-linker – PhoX; FDR – 1%; peptide and fragment tolerance – 20 ppm. Cysteine carbamidomethylation was set as a fixed modification, methionine oxidation, and asparagine deamidation were allowed as variable modifications. The resulting data were filtered for restraints present in all 3 replicates with a cumulative count of at least 7 cross-link matched spectra (CSMs).

Structural models used to visualize XL-MS results were generated using AlphaFold3.^58^ The models were predicted for MIB (111–749) together with trastuzumab IgG1 LC and HC (1–230) or CoV07 IgM LC and HC (1–338). The best-scoring models were compared with a previously reported antibody–MIB structure (Fig. S8).^28^ The XL-MS restraints were mapped on the models, analyzed, and visualized using ChimeraX 1.10.^59^

### High-speed atomic force microscopy

HS-AFM (RIBM, Japan) was conducted in tapping mode at RT in buffer, with free amplitudes of 1.5–2.5cnm and amplitude set points larger than 90%. Silicon nitride cantilevers with electron-beam deposited tips (USC-F1.2-k0.15, Nanoworld AG), nominal spring constants of 0.15cN/m, resonance frequencies around 500ckHz, and a quality factor of ∼2 in liquids were used. Antibody samples and MIB were combined at molar ratios of 1:2 for anti-WTA 4497 IgG1:MIB, 1:4 for SIgA:MIB, 1:10 for anti-WTA 4497 and anti-S-protein CoV07 IgM:MIB, and 1:12 for anti-CD52 campath IgG1-RGY:MIB at a final antibody concentration of 0.5 mg/mL and incubated at room temperature for 5 min. The working antibody concentration of 2 µg/mL was achieved via serial dilution using a weak immobilization buffer (75cmM NaCl, 10cmM Tris, 5cmM MgCl_2_, pHc7.4). Samples were immediately incubated on freshly cleaved muscovite mica for 40 s, followed by three washing steps using strong immobilization buffer (75cmM NaCl, 10cmM Tris, 10cmM NiCl_2_, pHc7.4). Imaging was performed in a strong immobilization buffer. Free antibodies of the respective variant were added to the sample after MIB-bound complexes image acquisition to facilitate direct comparisons between free and MIB-bound antibodies. Pure immunoglobulin samples were prepared as described above, using protein-free weak immobilization buffer instead of MIB.

Resulting HS-AFM images were processed using Gwyddion 2.62 and Fiji.^60–63^ For the analysis of Fab motion, HS-AFM movies were corrected for drift between subsequent frames via a slice alignment plugin based on image cross-correlation. Free and MIB-bound Fab trajectories were generated by employing a plugin for multiple particle detection and tracking (particle tracker 2D/3D). Trajectories (*r⃗*(*t*)) were further processed using in-house algorithms implemented in MATLAB (MathWorks). The mean square displacement *MSD*(Δ*t*) = 〈*r⃗*(*t*+ Δ*t*) - *r⃗*(*t*)〉^2^) was calculated as a function of the time-lag Δt between subsequent images. MSD(Δt) was further analyzed by employing a model for 2D diffusion with short-term mobility *D* within a circular confinement of radius R according to *MSD*(Δ*t*) = *R*^2^[1 - *exp*(- 4*D*Δ*t*/*R*^2^)].

## Acknowledgement

Recombinant IgG antibodies were a kind gift from Janine Schuurman and Aran Labrijn (Genmab, Utrecht, NL). Anti-S-protein CoV07 IgM and IgG1 were kindly provided by Paul Devine and Nick Bond, AstraZeneca (Cambridge, UK). The S-protein, S-protein RBD, and E1E2 (StrepTagII protein) were a kind gift from Laura Radic and Janke Schinkel (Amsterdam UMC, NL). Jack Li (Utrecht University) is acknowledged for providing the PhoX cross-linker. Furthermore, we would like to acknowledge members of the HeckLab (Utrecht University), namely: Sofia Kalaidopoulou Nteak for her suggestions regarding bottom-up proteomics data analysis, Linus Wollenweber for providing the polyclonal IgG sample, and Cynthia Kelley and Elena Giaretta for their suggestions.

## Author contributions

Conceptualization, T.K. and A.J.R.H; data curation, T.K., A.B.B., and J.S.; formal analysis, T.K., A.B.B., J.S., and J.P.; funding acquisition, A.J.R.H.; investigation, T.K., A.B.B., J.S. and J.P.; methodology C.d.H., S.H.M.R, M.N., R.L.; resources C.d.H., S.H.M.R, M.N., R.L.; supervision, S.H.M.R, J.P., and A.J.R.H.; visualization T.K., A.B.B., J.S., and J.P.; writing – original draft T.K., A.B.B., and A.J.R.H.; writing – review and editing, all authors.

## Data availability

Calibrated events for all MP measurements were deposited on Figshare. DIA LC-MS/MS and XL-MS data have been deposited to the ProteomeXchange Consortium via the PRIDE^64^ partner repository with the dataset identifier PXD078940.

