## Supplementary material for "Mycoplasma immunoglobulin binding protein universally binds human antibodies, thereby reducing Fab arm flexibility and displacing antigens in immune complexes": Suppl Figures and Tables

### List of contents:

#### Supplementary figures


1. Mass photometry of serum-derived IgG, IgA, IgM, and recombinant MIB.
2. Mass photometry analysis of polyclonal IgA and IgM binding to MIB.
3. Mapping interactions of monomeric antibodies with MIB.
4. Analyzing interactions of multimeric antibodies with MIB.
5. MIB binding consistently stabilizes antibodies across antibody isotypes.
6. HS-AFM images of unbound MIB, IgG, and IgM particles.
7. Inter- and intra-links detected in XL-MS experiment of antibody–MIB complexes and unbound MIB.
8. Comparison of structural models with MIB structure and XL-MS results.
9. MIB interferes with antibody–antigen complexes.
10. MIB forms complexes with IgG1 hexamers in absence and presence of C1q.
11. Reducing SDS-PAGE optimization of the cross-linker concentration for XL-MS.


#### Supplementary tables


1. Expected and measured masses in mass photometry experiments.
2. Overview of monoclonal antibodies and their complexes with MIB.
3. Heights and occupancies from the HS-AFM experiment.
4. Fab motion analysis.


#### Supplementary movies


1. HS-AFM movie of IgM-4497 recorded at 1 s/ frame and 2 nm/pixel played at 3 frames/s; z-scale 0–6.5 nm.
2. HS-AFM movie of IgM-4497 + MIB recorded at 1 s/frame and 2 nm/pixel played at 3 frames/s; z-scale 0–9 nm.
3. HS-AFM movie of IgG1-RGY with and without partial occupation by MIB recorded at 2 s/frame and 2 nm/pixel played at 3 frames/s; z-scale 0–8 nm.


#### Supplementary files


1. Filtered DIANN results table.
2. Median protein intensities and antibody annotation from the DIA LC-MS/MS quantification.
3. Filtered XL-MS pLink 3.0 results.

### Supplementary figures

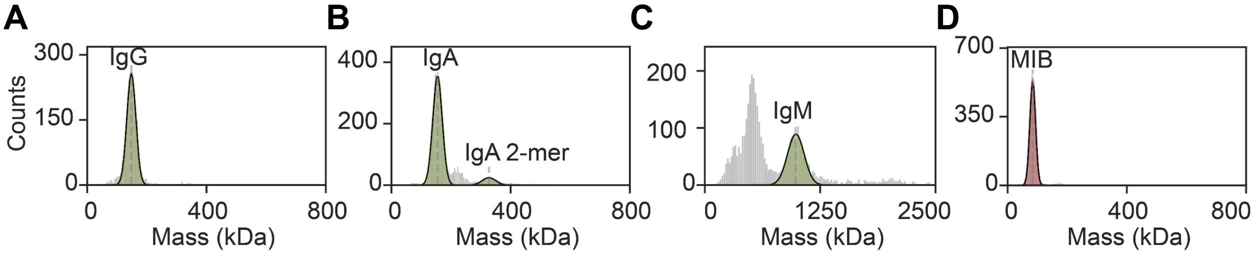


**Supplementary figure 1 Mass photometry of serum-derived IgG, IgA, IgM, and recombinant MIB.** Mass histograms of polyclonal antibodies: **A** IgG (147 ± 14 kDa), **B** IgA (154 ± 15 kDa and 327 ± 22 kDa), and **C** IgM (986 ± 89 kDa) and recombinant MIB (84 ± 9 kDa). The low-MW species observed in the polyclonal IgM sample correspond to previously observed co-purified contaminants.[1]

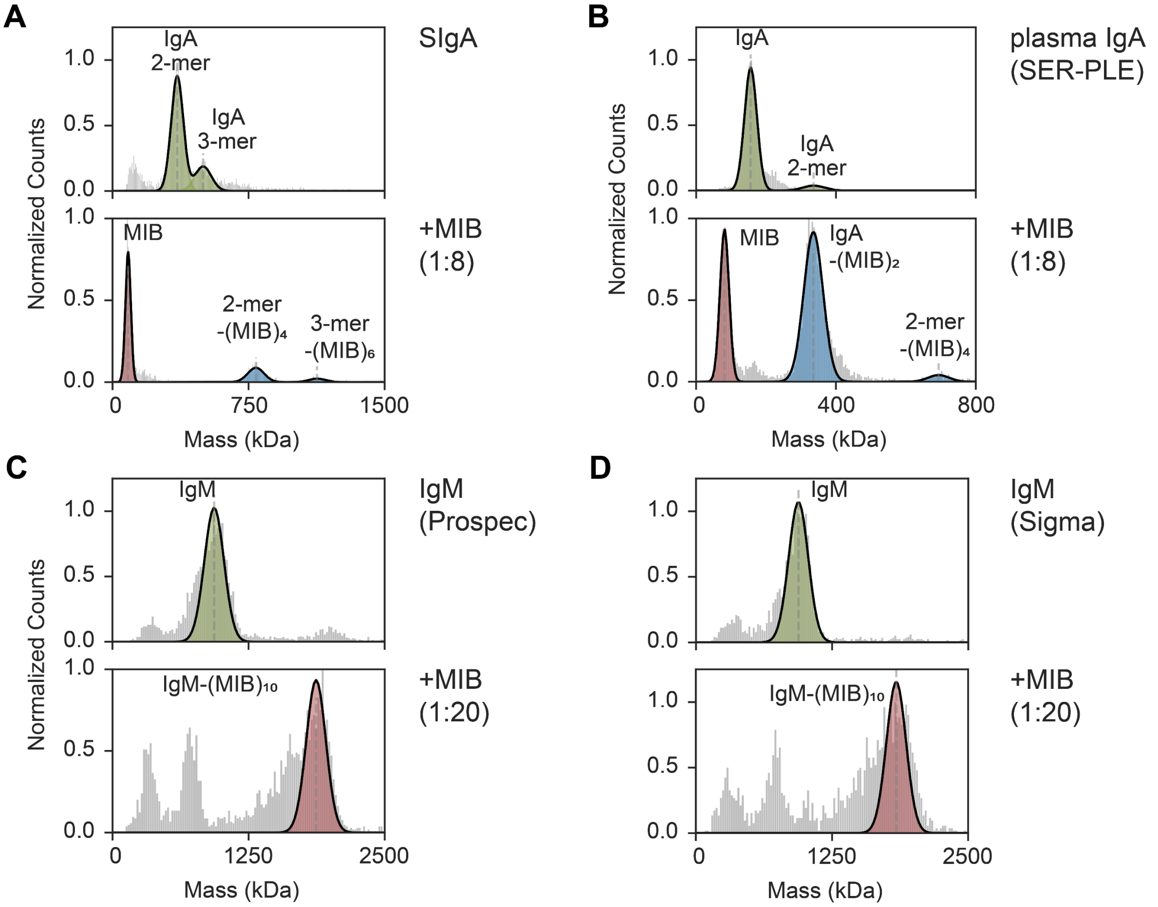


**Supplementary figure 2 Mass photometry analysis of polyclonal IgA and IgM binding to MIB.** Normalized mass histograms of: **A** secretory SIgA purified from human colostrum containing J-chain–coupled dimers and trimers (top) that bind 4 and 6 copies of MIB, respectively (bottom); **B** human plasma-isolated IgA containing monomers and a small portion of J-chain–coupled dimers (top), that bind 2 and 4 copies of MIB, respectively (bottom); commercially acquired IgM from **C** Prospec (ProSpec, Israel) and **D** Sigma-Aldrich (Merck, Germany), unbound control (top) and in complex with MIB (bottom). The results reveal that all studied antibodies reach full occupation of 2 MIBs/protomer. The annotated species correspond to following masses: **A** IgA dimer (356 ± 33 kDa), IgA trimer (500 ± 45 kDa); MIB (86 ± 13 kDa), IgA dimer+4MIB (791 ± 45 kDa), IgA trimer+6MIB (1127 ± 45 kDa); **B** IgA (156 ± 18 kDa), IgA dimer (336 ± 30 kDa); MIB (81 ± 13 kDa), IgA+2MIB (335 ± 28 kDa), IgA dimer+2MIB ( 693 ± 30 kDa); **C** IgM (934 ± 90 kDa); IgM+10MIB (1870 ± 90 kDa); **D** IgM (941 ± 90 kDa); IgM+10MIB (1840 ± 90 kDa).

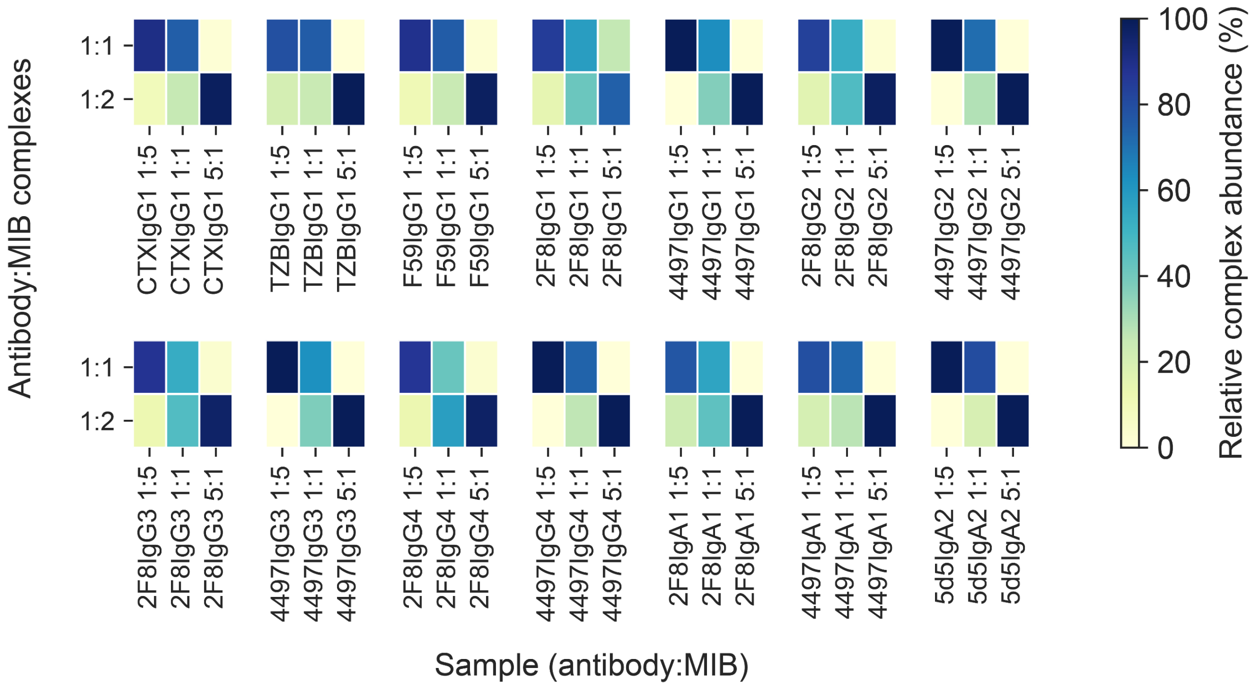


**Supplementary figure 3 Mapping interactions of monomeric antibodies with MIB.** Heatmap depicting relative abundances of antibody–MIB complexes for various IgG1-4 and IgA1-2 samples. To cover different aspects of antibody variability, we assessed: Fab-glycosylated cetuximab (CTX), κ-light chain variant, trastuzumab (TZB), and λ-light chain variant, F59. For two antibody variants, anti-WTA 4497 and anti-EGFR 2F8, we analyzed MIB binding across IgG1-4 and IgA1 subclasses. Finally, 5d5 IgA2 binding was assessed. Cumulatively, the data indicate no major differences in MIB interaction, either among antibodies of the same subclass recognizing different epitopes or between antibody isotypes.

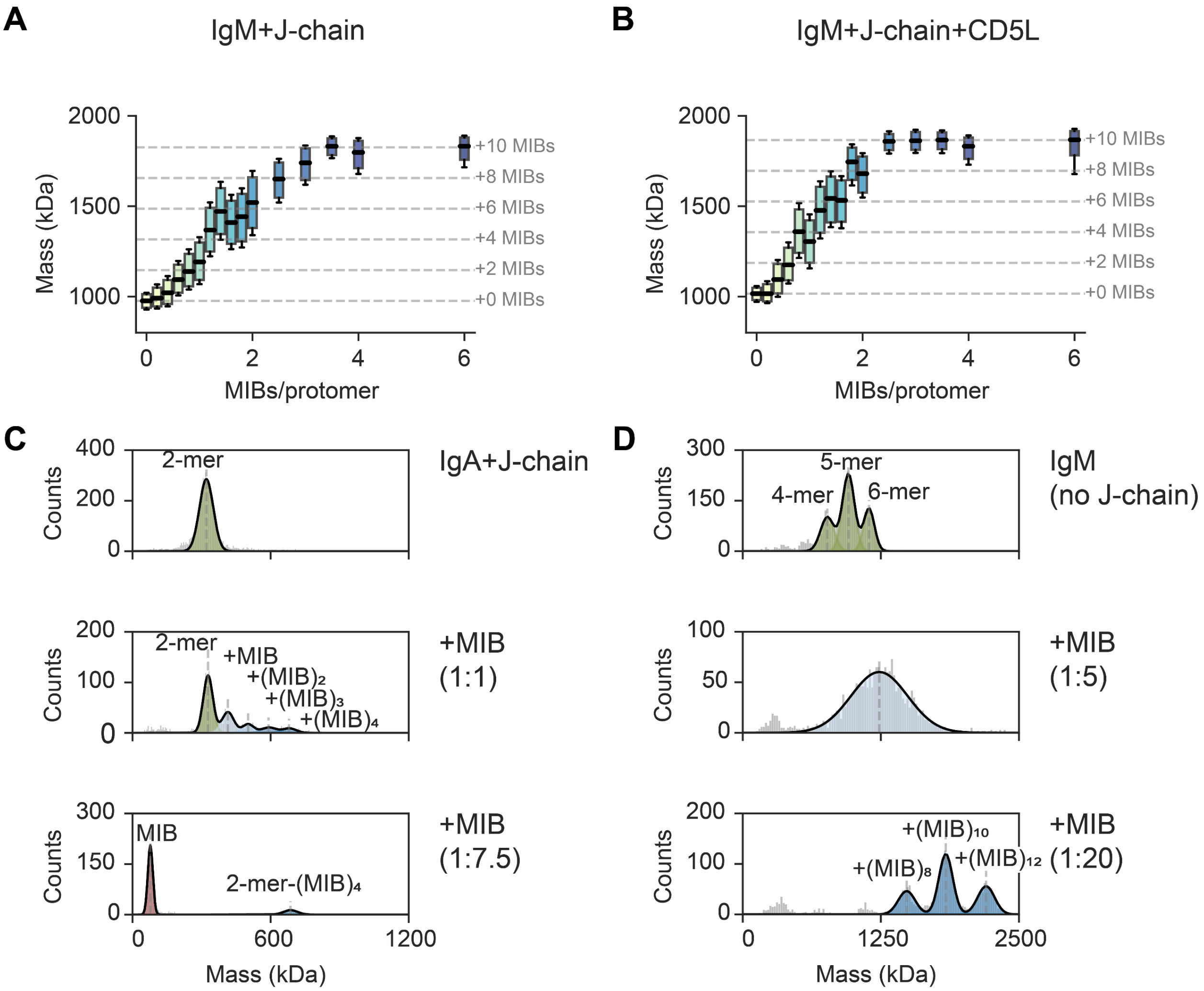


**Supplementary figure 4 Interactions of multimeric antibodies with MIB.** Box plots visualizing mass distributions obtained by mass photometry for **A** IgM+J-chain and **B** IgM+J-chain+CD5L, incubated with MIB in a range from 0 to 6 MIBs/protomer. The graphs show detected masses (boxes represent quartiles of mass distribution) on the y-axis and MIBs/protomer on the x-axis. The dashed lines represent the mass shifts corresponding to the binding of 2-10 MIBs (85 kDa) to the multimeric IgM antibody. Mass histograms visualizing **C** anti-WTA 4497 IgA+J-chain and **D** anti-WTA 4497 IgM (without J-chain) incubated with MIB. All mass photometry results show gradual occupation of antibodies upon titration by MIB, until reaching occupancy of 2 MIBs/protomer. Annotated species correspond to the following masses: **C** J-chain coupled IgA dimer (320 ± 30 kDa); J-chain coupled IgA dimer (328 ± 21 kDa), J-chain coupled IgA dimer + MIB (414 ± 26 kDa), J-chain coupled IgA dimer + 2MIB (502 ± 24 kDa), J-chain coupled IgA dimer + 3MIB (590 ± 30 kDa), J-chain coupled IgA dimer + MIB (680 ± 30 kDa); MIB (77 ± 11 kDa), J-chain coupled IgA dimer + 4MIB (686 ± 30 kDa), and **D** IgM tetramer (765 ± 61 kDa), IgM pentamer (956 ± 55 kDa), IgM hexamer (2242 ± 49 kDa); unresolved IgM-MIB complexes (1236 ± 256 kDa); IgM tetramer + 8MIB (1483 ± 76 kDa), IgM pentamer + 10MIB (1838 ± 63 kDa), IgM hexamer + 12MIB (2202 ± 77 kDa).

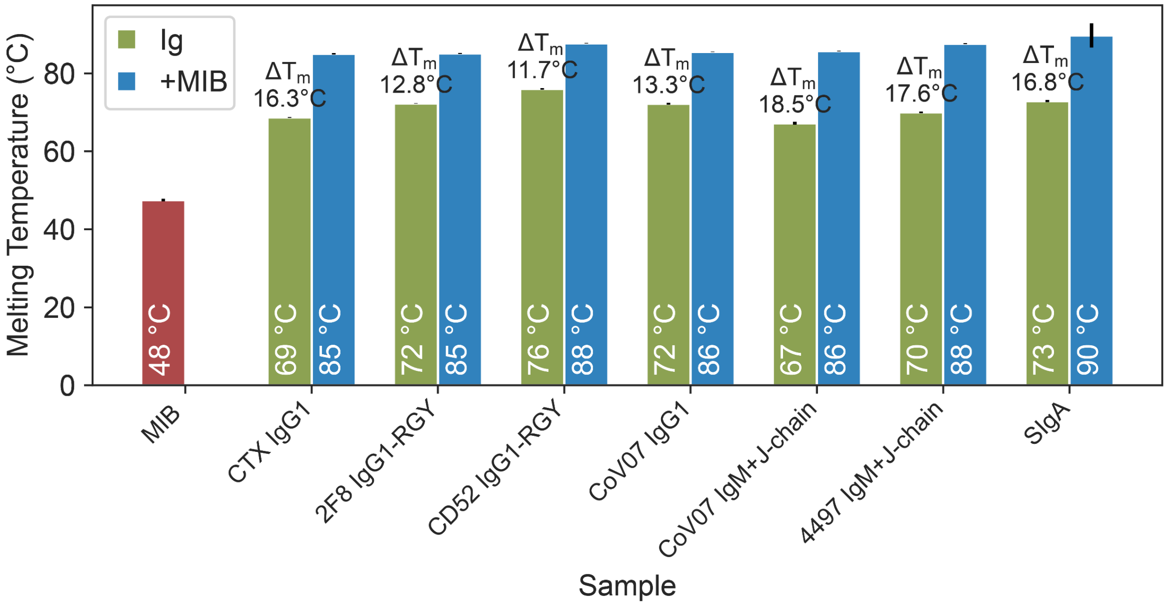


**Supplementary figure 5 MIB binding consistently stabilizes antibodies across antibody isotypes.** Differential scanning fluorimetry melting temperature determination for MIB (red), unbound antibodies (green), and antibody–MIB complexes (blue). The data were acquired for samples containing 2 MIBs per antibody protomer with a total protein concentration of 0.1 g/L in PBS, pH 7.4. The melting experiment was performed in n=3 replicates on Prometheus Panta (NanoTemper, Germany) in a temperature range of 25–95 ºC with a heating rate of 1 ºC/min.

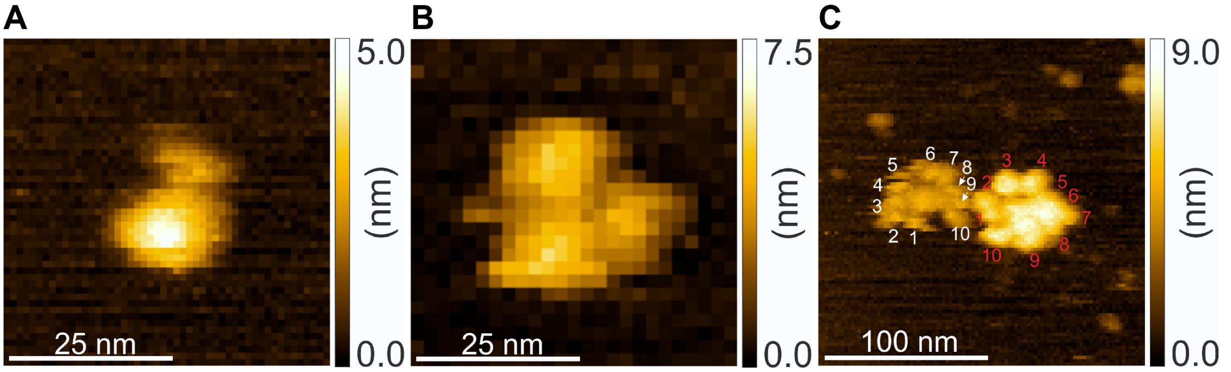


**Supplementary figure 6 HS-AFM images of unbound MIB, IgG, and IgM particles. A** unbound MIB, **B** unbound anti-WTA 4497 IgG1, and **C** anti-WTA 4497 IgM with MIB electrostatically immobilized on a mica surface. Panel **C** demonstrates the difference between MIB-bound IgM (right) and unbound IgM (left). It highlights that while the particle diameter remains unchanged, the height of MIB-bound IgM increases from approximately 4.4 nm for unbound IgM to 7.2 nm. Each Fab arm is annotated to facilitate the comparison.

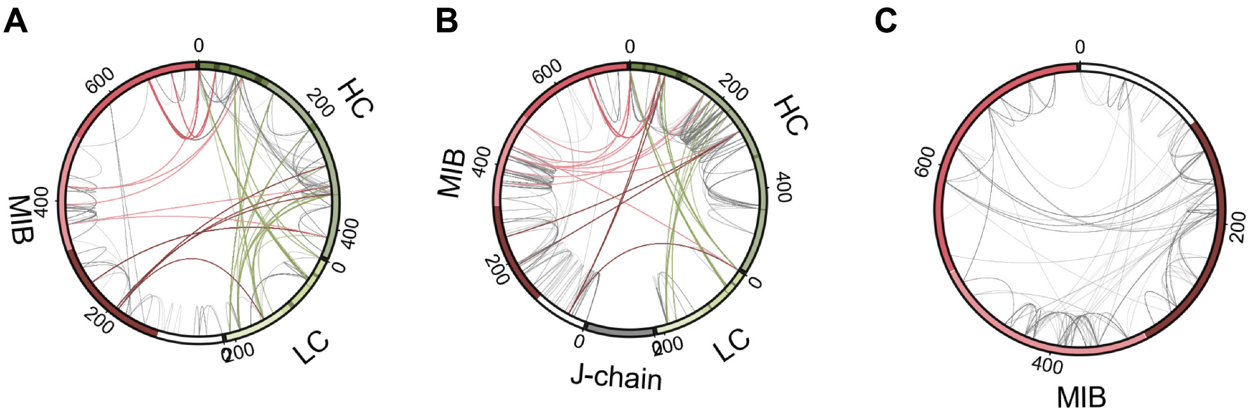


**Supplementary figure 7 Inter- and intra-links detected in XL-MS experiment of antibody–MIB complexes and unbound MIB.** Circos plots visualizing the XL-MS results for **A** trastuzumab IgG1–MIB, **B** CoV07 IgM–MIB, and **C** unbound MIB. The plot shows all cross-links detected in all n=3 replicates with at least 7 CSMs (cumulative count). All intra-links are shown in grey. The colors match the annotations in Fig. 4.

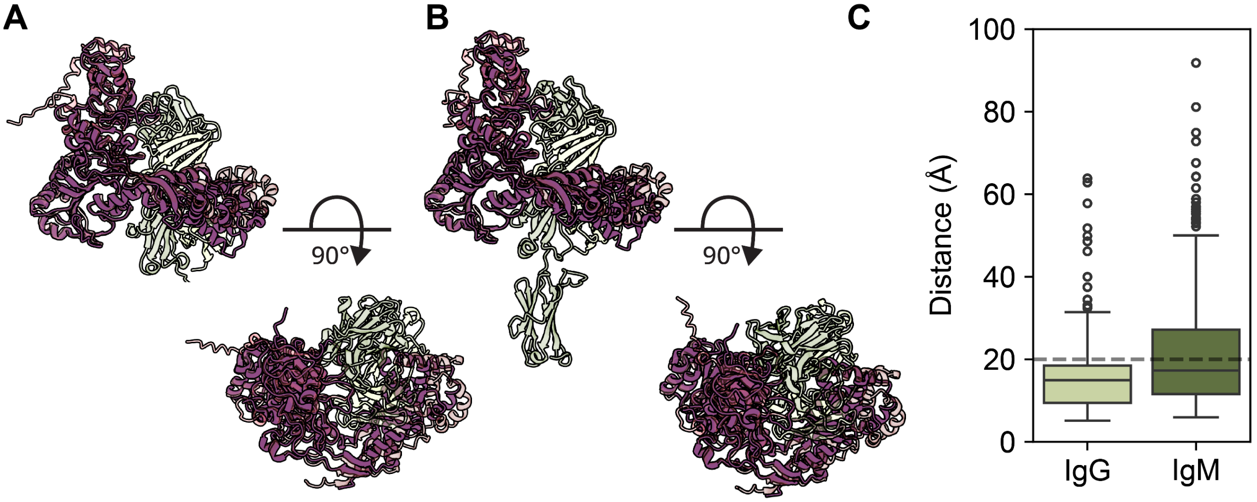


**Supplementary figure 8 Comparison of structural models with MIB structure and XL-MS results.** MIB structure (7ADM, purple)[2] overlaid with structural models of **A** trastuzumab IgG1–MIB and **B** CoV07 IgM–MIB complexes (semi-transparent). The comparison was performed in ChimeraX using Needleman-Wunsch algorithm.[3] **A** IgG model comparison to MIB (7ADM) yielded RMSD between 360 pruned atom pairs of 1.032 Å and 5.187 Å across all 600 pairs. **B** IgM model comparison to MIB (7ADM) yielded RMSD between 370 pruned atom pairs of 1.046 Å and 4.814 Å across all 600 pairs. **C** Distance distribution of all detected cross-links visualized on the structural models for IgG (light green) and IgM (dark green). The 20 Å distance cut-off [4] is visualized by a dashed line and suggests that the majority of restraints are satisfied. The over-length cross-links in the IgM sample are largely involving the CH1 domain and suggest a more compact conformation of the IgM-MIB complex, in line with the HS-AFM results.

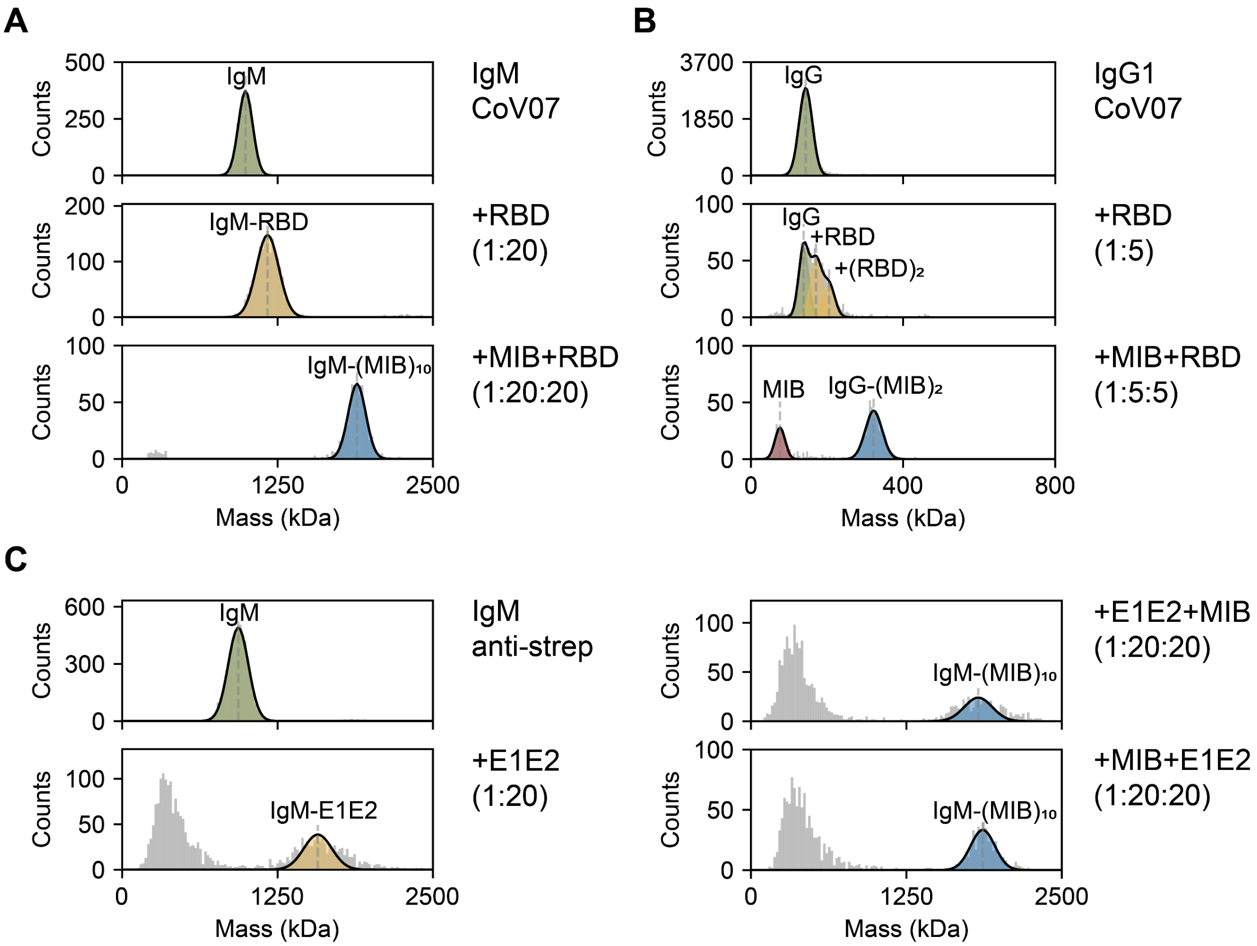


**Supplementary figure 9 MIB interferes with antigen-antibody complexes.** Mass photometry histograms depicting antibody–antigen interactions disrupted and obstructed by MIB. **A** CoV07 IgM+J-chain and **B** CoV07 IgG1 complexes with respective antigens are obstructed by MIB. Panels **A**, **B** show antibody control (top), antibody–antigen complex (middle), and antibody sample pre-incubated with MIB followed by antigen addition (bottom), highlighting that pre-incubation with MIB (5 min, RT) obstructs antibody–antigen complex formation**. C** anti-StrepTagII IgM+J-chain and its interaction with StrepTagII-containing protein E1E2 (88 kDa). The panel shows unbound IgM (top left), antigen-bound IgM (bottom left), disruption of antibody–antigen complex by MIB addition (top right), and obstruction of antibody–antigen complex formation by pre-incubating the antibody with MIB (bottom right). Annotated species correspond to following masses: **A** IgM (992 ± 58 kDa); IgM + RBD (1170 ± 90 kDa); IgM + MIB (1889 ± 71 kDa), **B** IgG1 (144 ± 17 kDa); IgG1 + RBD (171 ± 15 kDa), IgG1 + 2RBD (205 ± 15 kDa); MIB (76 ± 13 kDa), IgG1 + 2MIB (322 ± 22 kDa), and **C** IgM (934 ± 81 kDa); IgM + E1E2 (1573 ± 113 kDa); IgM + MIB (1827 ± 113 kDa); IgM + MIB (1864 ± 97 kDa).

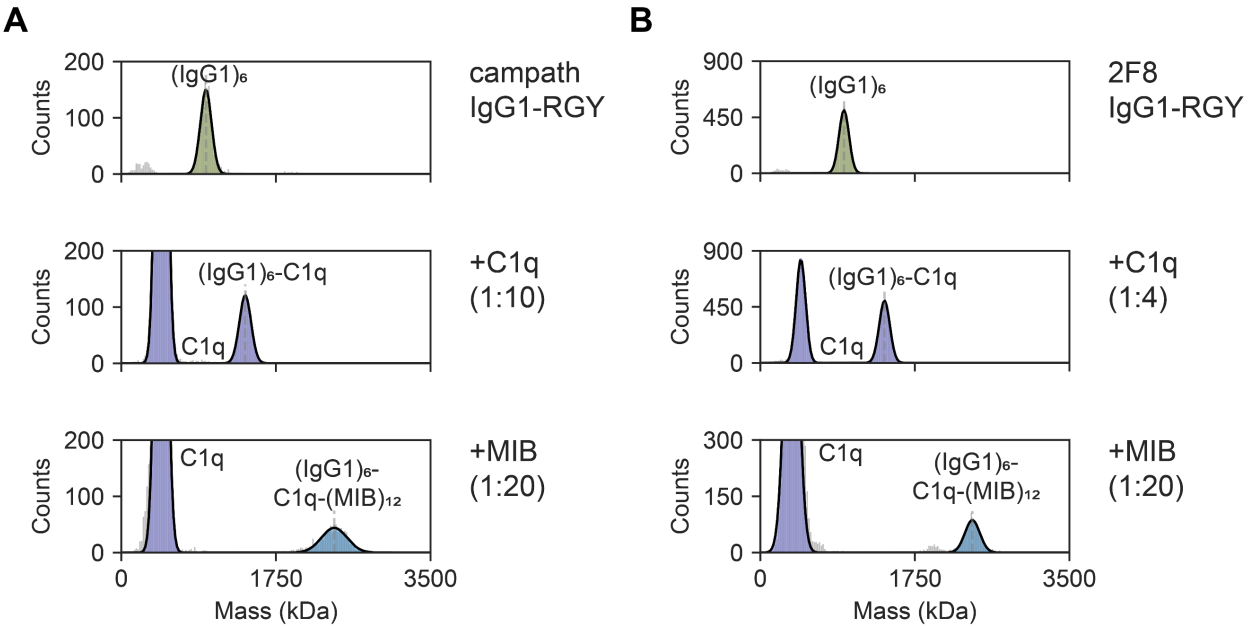


**Supplementary figure 10 MIB forms complexes with IgG1 hexamers in the absence and presence of C1q.** Mass histograms depicting interaction of **A** campath CD52 IgG1-RGY and **B** 2F8 IgG1-RGY with C1q (middle), and C1q and MIB (bottom). Annotated species correspond to following masses: **A** IgG1 hexamer (958 ± 64 kDa); C1q (453 ± 54 kDa), IgG1 hexamer+C1q (1402 ± 66 kDa); C1q (354 ± 75 kDa); IgG1 hexamer+C1q+12MIB (2401 ± 84 kDa), and **B** IgG1 hexamer (947 ± 57 kDa); C1q (459 ± 53 kDa), IgG1 hexamer+C1q (450 ± 60 kDa); C1q (451 ± 59 kDa); IgG1 hexamer+C1q+12MIB (2410 ± 144 kDa).

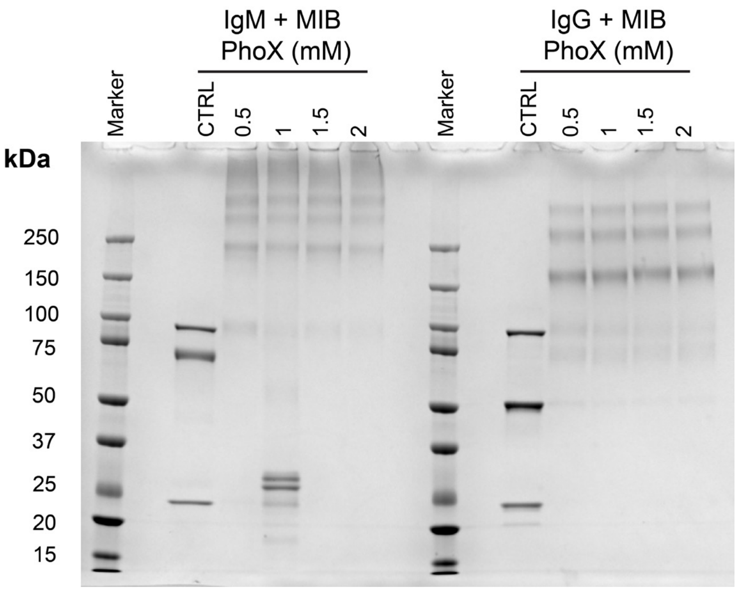


**Supplementary figure 11 Reducing SDS-PAGE optimization of the cross-linker concentration for XL-MS.** The experiment was performed using Criterion XT Bis-Tris 4 to 12% Precast gel, Criterion XT electrophoretic cell, XT Sample Buffer, XT MOPS Running Buffer, and Precision PluS-protein Dual Color Standard (Bio-Rad, USA), following the vendor-provided protocol. Each line contains 2 ug of protein. The gel was stained with Imperial Protein Stain (Thermo Scientific, USA).

### Supplementary tables

**Supplementary table 1 Expected and measured masses in mass photometry experiments.** The table is divided according to figures and panels in the main text. Expected masses were reported previously: IgG, IgG1, and $(IgG1)_6$[5], monomeric IgA, dimeric IgA+J-chain[6], serum-derived IgM+J-chain+CD5L and recombinant IgM+J-chain[1], S-protein[7], or calculated using ProtParam.[8]

| Figure | Panel | Sample | Annotated species | Expected mass (kDa) | Measured mass (kDa) |
|---|---|---|---|---|---|
| 2 | C | serum IgG (left) | IgG | 148 | 145 ± 15 |
| | | | IgG-(MIB) | 233 | 237 ± 18 |
| | | | IgG-$(MIB)_2$ | 318 | 330 ± 23 |
| | | | MIB | 85 | 78 ± 12 |
| | | | IgG-$(MIB)_2$ | 318 | 334 ± 23 |
| | | serum IgA (middle) | IgA | 158 | 151 ± 15 |
| | | | IgA-(MIB) | 243 | 236 ± 22 |
| | | | IgA-$(MIB)_2$ | 328 | 322 ± 23 |
| | | | MIB | 85 | 80± 13 |
| | | | IgA-$(MIB)_2$ | 328 | 330 ± 23 |
| | | | IgA dimer+J-chain-$(MIB)_4$ | 671 | 685 ± 23 |
| | | serum IgM (right) | IgM-$(MIB)_{3-5}$ | 1326 | 1321 ± 230 |
| | | | IgM-$(MIB)_{10}$ | 1836 | 1870 ± 105 |
| 3 | B | IgM+MIB (top, sample containing 3MIBs/protomer) | IgM-$(MIB)_{10}$ | 1820 | 1880 ± 10 |
| | | IgA1+J-chain+MIB (middle, sample containing 3MIBs/protomer) | MIB | 85 | 82 ± 10 |
| | | | IgA+J-chain-$(MIB)_4$ | 693 | 701 ± 22 |
| | | IgG1+MIB (bottom, sample containing 3MIBs/protomer) | MIB | 85 | 81 ± 12 |
| | | | IgG1-(MIB) | 233 | 243 ± 15 |
| | | | IgG1-$(MIB)_2$ | 318 | 335 ± 18 |
| 6 | A | CoV07 IgG1+RBD (top) | IgG1 | 148 | 138 ± 13 |
| | | | IgG1-(RBD) | 180 | 171 ± 15 |
| | | | IgG1-$(RBD)_2$ | 212 | 205 ± 15 |
| | | CoV07 IgG1+RBD+MIB (bottom) | MIB | 85 | 75 ± 14 |
| | | | IgG1-$(MIB)_2$ | 318 | 321 ± 23 |
| | B | CoV07 IgM+S-protein (top) | S-protein | 475 | 484 ± 57 |
| | | | IgM-$S_3$ | 2395 | 2422 ± 90 |
| | | | IgM-$S_4$ | 2870 | 2881 ± 90 |
| | | CoV IgM+S-protein+MIB (bottom) | S-protein | 475 | 488 ± 53 |
| | | | IgM-$(MIB)_{10}$ | 1820 | 1907 ± 72 |
| | C | anti-CD52 IgG1-RGY (top) | $(IgG1)_6$ | 888 | 958 ± 64 |
| | | anti-CD52 IgG1-RGY+MIB (bottom) | $(IgG1)_6$-$(MIB)_{12}$ | 1908 | 2061 ± 86 |

**Supplementary table 2 Overview of antibody–MIB complexes analyzed by mass photometry.** Summary of antibody isotypes and subclasses with maximum detected occupancies and MIBs/protomer for each sample. The results demonstrate that all studied antibodies reached the same occupancy of 2 MIBs/protomer.

| | Sample | Figure | Additional information | Protomers/ molecule | Maximum occupancy | Maximum MIBs/protomer |
|---|---|---|---|---|---|---|
| IgG1 | Trastuzumab | S3 | Kappa light chain | 1 | 2 | 2 |
| | F59 | S3 | Lambda light chain | 1 | 2 | 2 |
| | Cetuximab | S3 | Fab glycosylation | 1 | 2 | 2 |
| | 2F8 | S3 | | 1 | 2 | 2 |
| | 4497 | 3, S3 | | 1 | 2 | 2 |
| | CoV07 | 6A, S9 | | 1 | 2 | 2 |
| | serum polyclonal IgG | 1C | Single donor serum | 1 | 2 | 2 |
| IgG2 | 2F8 | S3 | | 1 | 2 | 2 |
| | 4497 | S3 | | 1 | 2 | 2 |
| IgG3 | 2F8 | S3 | | 1 | 2 | 2 |
| | 4497 | S3 | | 1 | 2 | 2 |
| IgG4 | 2F8 | S3 | | 1 | 2 | 2 |
| | 4497 | S3 | | 1 | 2 | 2 |
| IgA1 | 2F8 | S3 | | 1 | 2 | 2 |
| | 4497 | S3 | | 1 | 2 | 2 |
| | 4497 + J chain | 3 | Dimeric construct | 2 | 4 | 2 |
| | Serum/plasma polyclonal IgA | 1C, S2B | Pooled serum and single donor plasma-derived samples | 1/2 | 2/4 | 2 |
| IgA2 | 5d5 | S3 | | 1 | 2 | 2 |
| SIgA | colostrum polyclonal | S2A | Commercial sample, Sigma-Aldrich | 1/2/3 | 4/6/8 | 2 |
| IgM | CoV07+J-chain | 6B, S9A | | 5 | 10 | 2 |
| | anti-strep+J-chain | S9C | | 5 | 10 | 2 |
| | 4497 (no J-chain) | S4D | | 4/5/6 | 8/10/12 | 2 |
| | 4497+J-chain | 3, S4A | | 5 | 10 | 2 |
| | 4497+J-chain+ CD5L | S4B | | 5 | 10 | 2 |
| | serum polyclonal (IgA+J-chain+pIgR) | 1C, S2C–D | Pooled serum and 2 commercial samples | 5 | 10 | 2 |
| IgG1 | anti-CD52 campath | S10A | | 6 | 12 | 2 |
| | anti-EGFR | S10B | | 6 | 12 | 2 |
| | anti-CD38 005 | 7A–B | | 6 | 12 | 2 |

**Supplementary table 3 Heights and occupancies observed in the HS-AFM experiment.** Heights and occupancies obtained for unbound antibodies and MIB-bound complexes. The estimated height was calculated as a sum of unbound Fab height and MIB height of 4.2±0.4 nm (n=21).

| Sample | Complex + MIB | | | Unbound antibody | | |
|---|---|---|---|---|---|---|
| | Fab | Fc | MIB | Fab | Fab + MIB | Fc |
| | Height (nm) | Height (nm) | Occupancy | Height (nm) | Estimated height (nm) | Height (nm) |
| **IgM anti-S-protein CoV07** | 7.1±0.3 (n=22) | 5.6±0.8 (n=16) | 9.2±2.1 (n=34) | 4.4±0.2 (n=17) | 8.6±0.5 | 5.2±0.3 (n=18) |
| **IgM anti-WTA 4497** | 7.2±0.3 (n=33) | 5.3±0.6 (n=22) | 9.8±0.4 (n=29) | 4.4±0.4 (n=29) | 8.6±0.6 | 5.1±0.3 (n=13) |
| **IgG1 anti-WTA 4497** | 7.1±0.8 (n=33) | 3.9±0.6 (n=16) | 1.9±0.3 (n=18) | 4.0±0.2 (n=20) | 8.2±0.5 | 3.6±0.5 (n=16) |
| **SIgA** | 7.2±0.4 (n=33) | 9.5±0.8 (n=21) | 3.8±0.5 (n=23) | 4.3±0.8 (n=31) | 8.5±0.9 | 8.1±1.7 (n=20) |

**Supplementary table 4 Fab motion analysis.** Mobilities (D, $nm^2/s$) obtained from linear fit analysis of the mean square displacement (MSD(Δt)) and mobilities (D, $nm^2/s$) and circular confinement of radius (R, nm) obtained from a 2D diffusion model analysis of MSD(Δt) for MIB-bound and unbound antibodies.

| Sample | | Linear fit | Confined diffusion | |
|---|---|---|---|---|
| | | D ($nm^2/s$) | D ($nm^2/s$) | R (nm) |
| **IgM anti-WTA 4497** | **unbound** | 2.99 ± 0.59 | 5.9 ± 0.7 | 10.3 ± 0.6 |
| | **+MIB** | 0.34 ± 0.19 | 0.56 ± 0.9 | 4.5 ± 0.4 |
| **IgG1-RGY anti-CD52 campath** | **unbound** | 1.24 ± 0.25 | 2.5 ± 0.2 | 8.7 ± 0.4 |
| | **+MIB** | 0.04 ± 0.04 | 0.55 ± 0.18 | 2.4 ± 0.2 |
